# Digital museum of retinal ganglion cells with dense anatomy and physiology

**DOI:** 10.1101/182758

**Authors:** J. Alexander Bae, Shang Mu, Jinseop S. Kim, Nicholas L. Turner, Ignacio Tartavull, Nico Kemnitz, Chris S. Jordan, Alex D. Norton, William M. Silversmith, Rachel Prentki, Marissa Sorek, Celia David, Devon L. Jones, Doug Bland, Amy L. R. Sterling, Jungman Park, Kevin L. Briggman, H. Sebastian Seung, the EyeWirers

**Author notes:** http://eyewire.org (the EyeWirers). Present addresses: Dept. of Structure and Function of Neural Networks, Korea Brain Research Institute, Daegu 41068, Republic of Korea (JSK); Center of Advanced European Studies and Research (caesar), Dept. of Computational Neuroethology, 53175 Bonn, Germany (KLB). Co-first authors (J. Alexander Bae, Shang Mu, Jinseop S. Kim, Nicholas L. Turner).

## Abstract

Most digital brain atlases have macroscopic resolution and are confined to a single imaging modality. Here we present a new kind of resource that combines dense maps of anatomy and physiology at cellular resolution. The resource encompasses almost 400 ganglion cells from a single patch of mouse retina, and a digital “museum” provides a 3D interactive view of each cell’s anatomy as well as graphs of its visual responses. To demonstrate the utility of the resource, we use it to divide the inner plexiform layer of the retina into four sublaminae defined by a purely anatomical principle of arbor segregation. We also test the hypothesis that the aggregate neurite density of a ganglion cell type should be approximately uniform (“density conservation”). Finally, we find that ganglion cells arborizing in the inner marginal sublamina of the inner plexiform layer exhibit significantly more sustained visual responses on average.

---

Calcium imaging followed by 3D electron microscopy (EM) has become established as a powerful approach for obtaining anatomical and physiological information about the same neurons. The approach has been used to study visual neurons in mouse cortex [Bock et al., 2011, Lee et al., 2016] and retina [Briggman et al., 2011], and oculomotor neurons in larval zebrafish hindbrain [Vishwanathan et al., 2017]. These prior studies have been limited to tens of neurons sparsely sampled from a single individual. Here we present as a resource the anatomy and physiology of almost 400 ganglion cells (GCs) densely sampled from a single patch of mouse retina.

To facilitate exploratory data analysis using the resource, we have constructed the Eyewire Museum (http://museum.eyewire.org), where every reconstructed ganglion cell can be interactively viewed along with its visual response properties. Due to its cellular and subcellular resolution, the Museum is novel relative to traditional brain atlases, which typically divide the brain into macroscopic regions [Lein et al., 2007, Amunts et al., 2013, Zingg et al., 2014]. Because it fuses anatomy with physiology, the Museum is also novel relative to traditional atlases of neuronal morphologies such as `neuromorpho.org` [Ascoli et al., 2007] and `wormatlas.org` [Hall et al., 2007]. Previous dense EM reconstructions in the mouse retina [Helmstaedter et al., 2013] and larval zebrafish olfactory bulb [Wanner et al., 2016] were limited to anatomy only. A recent large-scale calcium imaging study contains the visual responses of more than 11,000 neurons from 50 retinas, but less than 1% of the cells had their dendritic arbors reconstructed [Baden et al., 2016].

To illustrate the utility of our resource, we use it to reveal new principles of retinal organization. First, we show how to optimally subdivide the inner plexiform layer (IPL) of the retina using the purely anatomical principle that arbors should segregate into distinct sublamina. For GC dendritic arbors, segregation is maximized by subdividing the IPL into two marginal sublamina flanking a central sublamina. For bipolar cell (BC) axonal arbors, in contrast, segregation is maximized by subdividing the IPL into inner and outer sublamina. Using the marginal-central and inner-outer dichotomies as a starting point, we divide our dense sample of GCs into six high-level clusters, which are subdivided further to end up with 47 clusters. This raises the question of how to validate whether clusters are indeed GC types.

To answer this question, we propose a “density conservation” principle: the aggregate arbor density of a GC type should be approximately uniform across the retina. Our principle is meant to supersede the traditional principle that the dendritic arbors of a GC type should “tile” the retina with little overlap [Wässle et al., 1981]. The latter seems inconsistent with observations of substantial overlap between dendritic arbors for mouse GC types [Zhang et al., 2012, Rousso et al., 2016]. Density conservation is shown to be satisfied by 24 of our clusters, which are internally validated as pure types. Six of these types appear novel, in the sense that we have been unable to find any matching reports in the literature.

Using the calcium imaging data, we have the opportunity to relate the above structural analyses to retinal function. It was previously proposed that marginal GCs have more sustained responses to visual stimuli than central GCs [Roska and Werblin, 2001]. We reexamine this conventional wisdom, and confirm that inner marginal GCs are significantly more sustained on average, but other high-level GC clusters do not exhibit statistically significant differences in sustainedness. The analogous finding has recently been reported for mouse BCs [Franke et al., 2017].

## Dense anatomy and physiology

We present a large-scale survey of almost 400 GCs in the mouse retina, combining anatomical information from serial block face scanning electron microscopy [Denk and Horstmann, 2004] with physiological information from calcium imaging. The set of reconstructed GCs is dense, meaning that it includes the arbor of every soma inside a 0.3 × 0.35 mm^2^ patch of mouse retina. Visual responses from calcium imaging are available for 82% of the cells. Our survey is based on the e2198 dataset, a small fraction of which was previously used to study retinal circuits for motion computation [Briggman et al., 2011, Kim et al., 2014, Greene et al., 2016].

A previous dense sample of GCs from the e2006 dataset contained 10× fewer cells, lacked physiological information, and the arbors of all but the smallest cells were severely cut off by the borders of the (0.1 mm)^2^ patch of mouse retina [Helmstaedter et al., 2013]. In previous large-scale anatomical surveys using light microscopy, GCs were sparsely sampled from many retinas, and lacked physiological information [Badea and Nathans, 2004, Kong et al., 2005, Coombs et al., 2006, Völgyi et al., 2009, Sümbül et al., 2014].

As previously described in Briggman et al. [2011], the e2198 dataset had a physiological component (time series of visual responses observed via two-photon calcium imaging *ex vivo*) and an anatomical component (3D image stack from serial block face scanning electron microscopy). Both imaging techniques were targeted at the same patch of a single mouse retina (Fig. 1A). The orientation of the patch relative to body axes was inferred from the reconstructed GCs (Methods), as information about the orientation of the retina was not recorded at the time of dissection.

**Figure 1:**
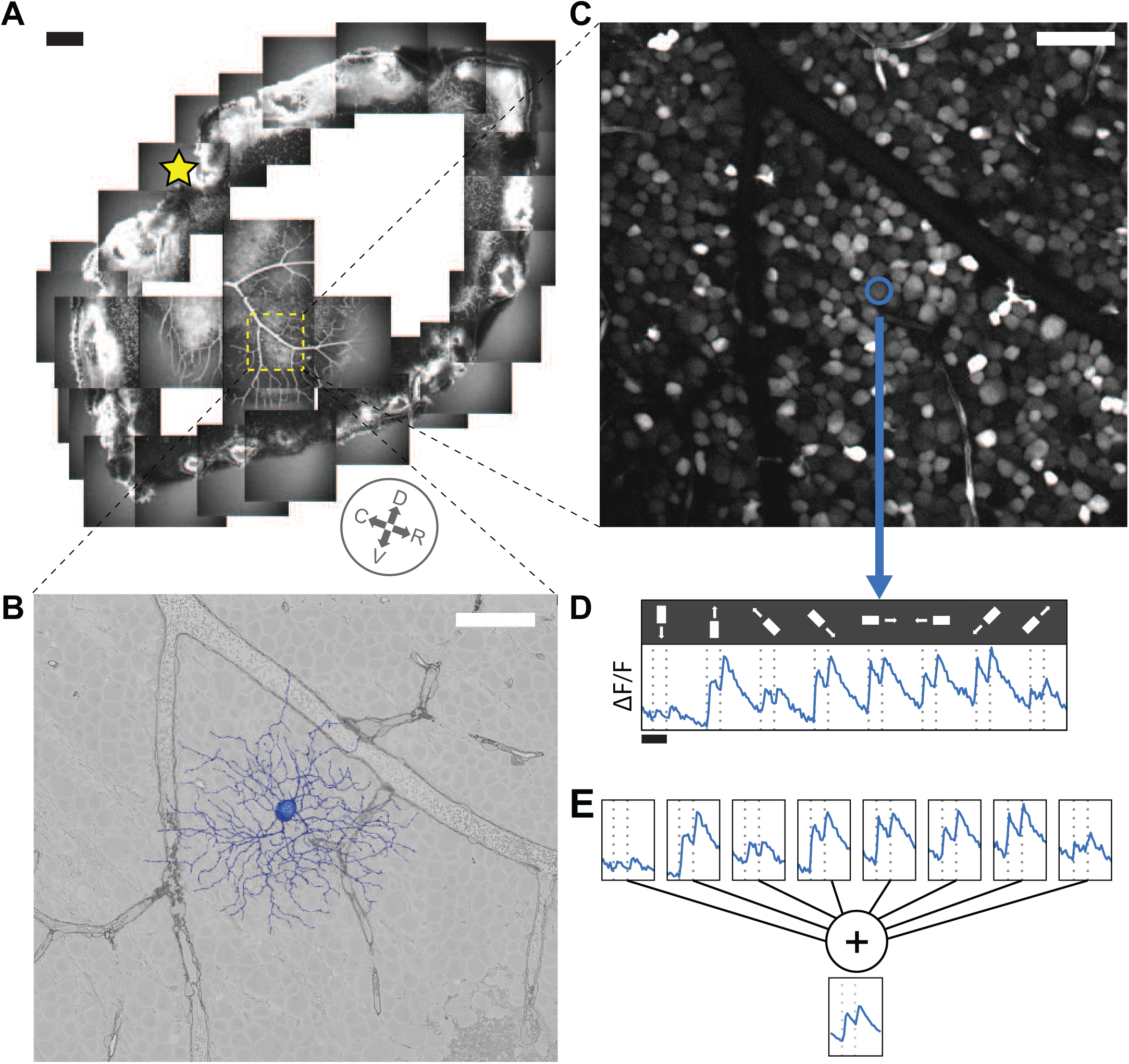
Anatomy and Physiology of Retinal Ganglion Cells via Electron and Light Microscopy. (**A**) Hemiretina containing imaged 0.3 × 0.35 mm^2^ patch (yellow square). Star, optic disk. Compass rosette, inferred cardinal directions (dorsal, ventral, rostral, caudal, see Methods). (**B**) 3D reconstruction of GC dendritic arbor (blue) and 2D cross section through GCL in serial EM image (grayscale). (**C**) Soma of same GC (blue) in image of GCL obtained via two-photon microscopy. (**D**) Fluorescence versus time for same GC along with stimulus sequence of light bar moving in eight directions on dark background (see main text for true aspect ratio of bar). (**E**) Averaging over stimulus directions (shown) and trials (not shown) yields temporal response function for GC. Scale bars, 200 *μ*m (A), 50 *μ*m (**B**, **C**) and 2 sec (**D**).

In the anatomical dataset (Fig. 1B), every cell body in the ganglion cell layer (GCL) was examined for evidence of an axon. Displaced GCs, with cell bodies in the inner nuclear layer (INL), were neglected by our survey. The axon emanated directly from the cell body in some cases, and branched from a primary dendrite in others. Through this systematic search, we identified a total of 396 GCs in e2198. Many of the GCs are rendered in Item S2 and the Movie S1. The dendritic arbors of all cells were reconstructed by almost 30,000 members of the online community known as Eyewire (Methods, Item S5). For subsequent analyses, we also made use of reconstructions of starburst amacrine cells (SACs) and bipolar cells (BCs) from previous studies [Kim et al., 2014, Greene et al., 2016].

Eleven cells were so severely cut off by the borders of the imaged volume that they were discarded. Four more cells were discarded as “weirdos” that may have resulted from developmental abnormalities. Excluding severely cut-off and “weirdo” cells left a sample of 381 that were used for subsequent analysis. All cells, including cut-off and “weirdo” cells, can be examined interactively at the Eyewire Museum.

Previous light microscopic surveys sparsely sampled GCs from many locations in many retinas [Badea and Nathans, 2004, Kong et al., 2005, Coombs et al., 2006, Völgyi et al., 2009, Sümbül et al., 2014]. The arbor diameter and density of alpha GCs are known to depend strongly on retinal location [Bleckert et al., 2014]. Asymmetry [Kim et al., 2008] and stratification depth [Sümbül et al., 2014] of the J cell also depend on retinal location. Because the cells of our dense sample come from a single location in a single retina, variation within a GC type is expected to be relatively small, facilitating accurate classification of GC types.

Some sparse sampling techniques used in the light microscopic surveys might miss GC types due to selection bias. Our dense GC sample is expected to include examples of almost all GC types. Possible exceptions would be types that are very rare and happen by chance not to occur in our finite sample, or types that are systematically absent in the region of the retina that contains the sample. Regarding the latter possibility, we are not aware of GC types that are known to exist in one retinal region but not another.

Our GC sample also differs from previous light microscopic surveys because it comes with visual responses from calcium imaging. In the physiological dataset (Fig. 1C), regions of interest (ROIs) were manually drawn around cell bodies in the GCL. For each ROI, the time series of the calcium signal was computed as the sum of the pixels in the ROI for each time point (Fig. 1D). We were able to extract calcium signals for 326 cells, 82% of the 396 GCs that were reconstructed from the anatomical dataset. The remaining cells lacked meaningful signals, either because they lacked sufficient calcium indicator, lacked sufficiently strong visual responses, or were too far from the focal plane used for two-photon imaging. Briggman et al. [2011] only reported calcium signals for 25 On-Off direction selective GCs, a small fraction of the current dataset.

The visual stimulus was a light bar on a dark background, with width 0.2 mm and length 1 mm (Fig. 1D). In each stimulus trial, the bar moved at 1 mm/sec along its long axis. Eight directions of movement were used to evaluate direction selectivity (DS). Two-photon imaging was sequentially applied to each tile in a 3 × 3 array that covered the retinal patch with slight overlap. The vertical lines in Fig. 1D indicate when the leading and trailing edges of the bar crossed the center of the imaged tile. These nominal stimulus onset and offset times differ slightly from the true times, because the receptive fields of cells in the imaged tile vary in their location and size.

## Digital museum

Early neuroanatomists relied heavily on qualitative descriptions with text and drawings [Cajal, 1893]. Since then neuroanatomists have increasingly adopted quantitative and computational methods, yet visualization remains important for the exploratory stage of data analysis. We have constructed the Eyewire Museum to enable interactive visualization of 3D reconstructions of the GCs. A major novelty is the well-developed functionality for viewing any subset of neurons of the dataset in combination, unlike previous digital atlases that show only one neuron at a time [Ascoli et al., 2007, Hall et al., 2007]. Simultaneous display of cells enables visualization of overlap between arbors, which indicates potential for connectivity [Masland, 2004, Stepanyants and Chklovskii, 2005]. Every neuron in the dataset has a unique ID number. Entering a list of cell IDs into the search bar results in a 3D rendering of the cells displayed together, and a sharable URL containing the cell IDs (Fig. 2B). For fast alterations of the displayed subset of cells, the right collapsible sidebar contains a list of the selected cell IDs. Mousing over a cell ID causes that individual cell to be highlighted (Fig. 2I). Clicking on a cell ID makes it vanish. All of the above visualization features are useful for exploring the spatial relationships between cells, or equivalently for viewing a cell in the context defined by other cells.

**Figure 2:**
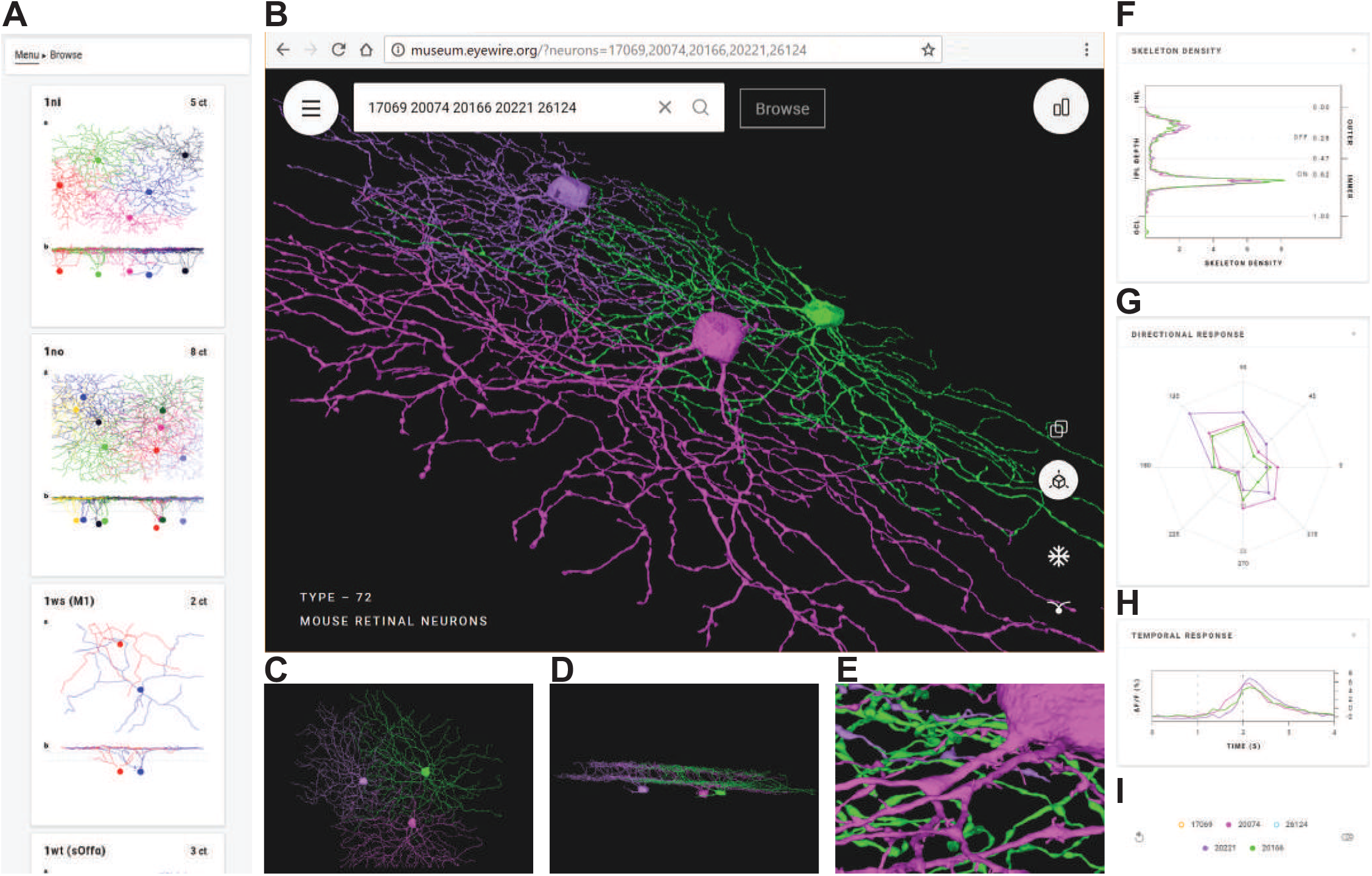
Eyewire Museum (`museum.eyewire.org`). (**A**) Left collapsible sidebar enables browsing GC clusters. (**B**) Display of 3D rendering of selected cells. URL containing the cell IDs can be shared. (*C*) View along the light axis (“Whole mount” view) of the retina. (*D*) View along the tangential axis. (*E*) Zoomed-in view. (**F-H**) Anatomical and physiological properties included in right collapsible sidebar; stratification profiles (**F**), directional response (**G**), and temporal response (**H**). (**I**) Right collapsible sidebar shows a list of selected cells. Mousing over a cell ID highlights the cell and its properties and cells can be removed from the display.

Visualizing a cell from different viewpoints is important for gaining different kinds of information. The orientation of the displayed cells can be manipulated arbitrarily using the mouse. Two specific viewpoints can be chosen by clicking on icons. The view along the light axis is the traditional “whole mount” view of the retina, and is best for visualizing properties such as dendritic path length, number of branch points, and so on (Fig. 2C). The view along a tangential axis reveals the depth at which arbors stratify in the IPL (Fig. 2D). Stratification depth is one of the most important features for clustering retinal neurons into cell types, as we will see later. One can also choose between orthographic and perspective views. The former is helpful for accurate perception of stratification depth when viewing along a tangential axis, while the latter gives a stronger 3D percept.

Neuroanatomy is multiscale, in the sense that structural properties at large and small length scales may be combined for scientific discovery. The Museum allows zooming out to visualize the overall shape of a cell’s arbor, and zooming in to see varicosities and other fine anatomical features (Fig. 2E). Rich multiscale functionality is made possible by the use of true 3D reconstructions, rather than skeleton representations of cells.

Graphs of anatomical and physiological properties are also available in the right collapsible sidebar (Fig. 2F, G, H). The first graph is the “stratification profile,” defined as the linear density of arbor as a function of IPL depth (Fig. 2F, Methods). The profile is treated like a probability distribution, with its area normalized to unity. The second graph shows the response of a cell as a function of stimulus direction (Fig. 2G). The third graph shows the “temporal response function” of a cell, defined as visual response versus time, averaged over trials and directions of the moving bar (Fig. 2H). By default the graphs show plots for all selected cell IDs (Fig. 2I). Mousing over the list of selected cell IDs singles out the plot for that individual cell in all three graphs.

While viewing arbitrary groups of cells is possible, the Museum contains predefined groups which will most commonly be viewed. One can browse through the groups using a left collapsible sidebar (Fig. 2A). The predefined groups were found by the hierarchical clustering procedure to be described later on. Some groups have been validated as neuronal cell types by procedures that will be described later.

The design of the Museum was inspired by our large, dense sample of retinal ganglion cells, which is currently a rare kind of dataset. As 3D EM becomes more widespread, however, we expect that many other researchers will want to build similar digital museums to promote exploration of large-scale reconstructions. In anticipation of their needs, we have made the Museum code publicly available.

We found the Museum visualizations complementary to our quantitative analyses of the dataset, which have uncovered several organizing principles for the IPL. We turn now to the first of these analyses concerning IPL structure along the light axis.

## Arbor segregation principle

In his description of ganglion cell diversity, Cajal [1893] divided the IPL into five sublaminae (S1-S5). It is unclear whether the five-way division is a subjective convention or has objective meaning. Wässle [2004] proposed a molecular basis (calbindin or calretinin staining) for the three borders between S1, S2, S3, and S4/5. Again, it is unclear whether the stained bands are more than arbitrary conventions. Famiglietti and Kolb [1976] divided the IPL into sublamina *a*, specialized for processing of Off (dark) stimuli, and sublamina *b*, specialized for processing of On (light) stimuli. This two-way division has objective meaning, but is rather coarse.

Here we show that the IPL can be divided into sublaminae based on the purely anatomical principle that BC arbors should segregate in the sublaminae. We normalize IPL depth so that it ranges from 0 to 1, where 0 denotes the border with the INL and 1 the border with the GCL (Methods). Suppose that we divide the IPL into inner and outer sublaminae. The inner-outer boundary is located at 0.47 IPL depth for now; the optimality of this value will be demonstrated later. A histogram of the difference between the amounts of inner arbor and outer arbor reveals that BCs separate into two clusters (Fig. 3A). Cells of one cluster have a mostly inner arbor, while cells of the other cluster have a mostly outer arbor (Fig. 3B). The gap in the center of the histogram indicates rarity of cells that evenly straddle the inner-outer boundary (Fig. 3A).

**Figure 3:**
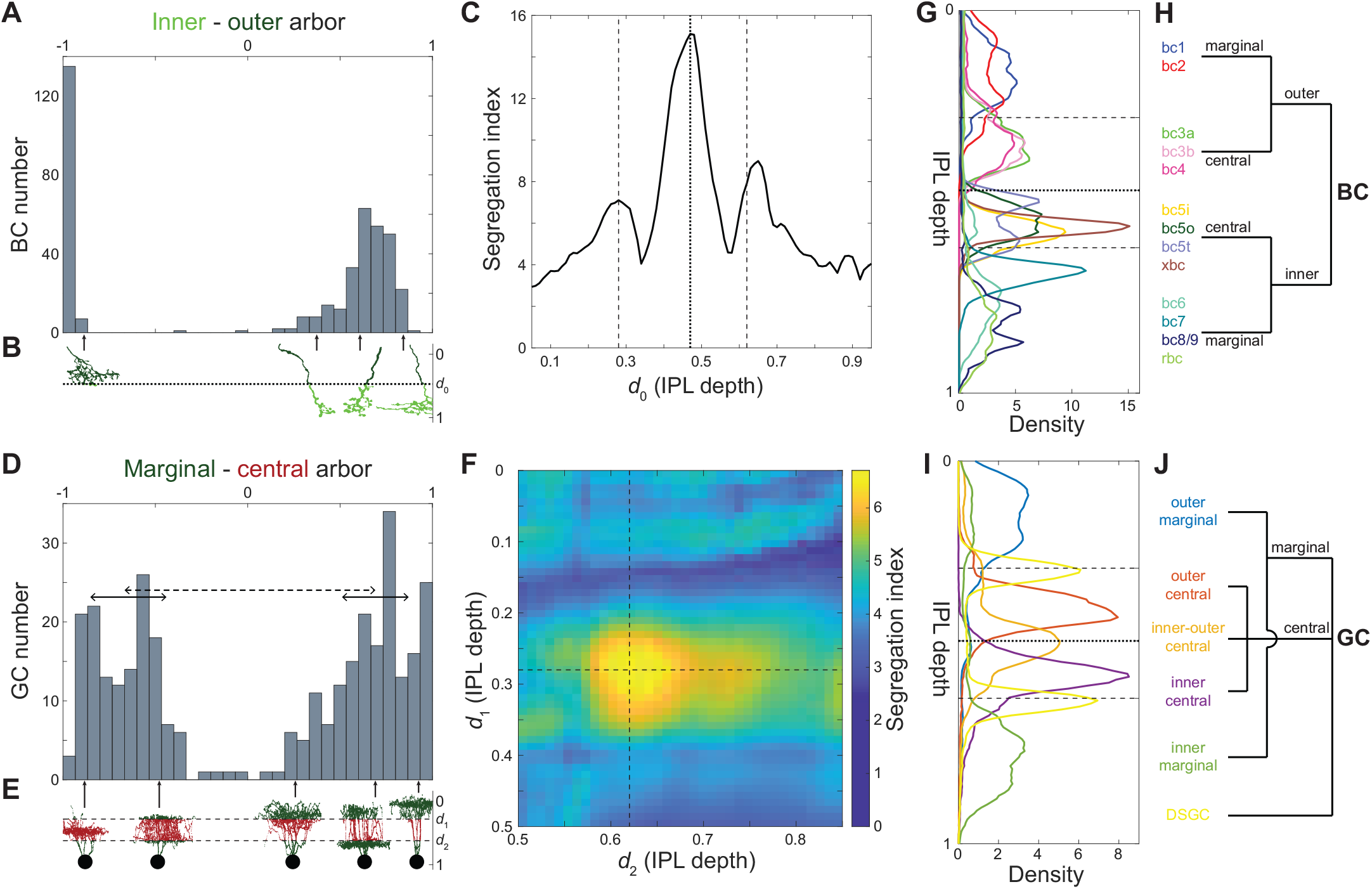
Maximizing Laminar Segregation of Arbors Yields Marginal-Central and Inner-Outer Divisions of the IPL. (**A**) Histogram of the difference between inner and outer arbor volume for BCs (total volume normalized to one). BC axonal arbors are either mostly inner (right cluster) or mostly outer (left cluster); intermediate cases are rare. (**B**) Inner arbor (light green) and outer arbor (dark green) of example BCs. The depth of the inner-outer boundary is denoted by *d*_0_. (**C**) BC inner-outer segregation is maximized for *d*_0_ *=* 0.47 (dotted line, same value used in **A**), with two flanking local maxima at or near the SAC depths (dashed lines). (**D**) Histogram of the difference between marginal and central arbor length for GCs (total length normalized to one). GC dendritic arbors are either mostly marginal (right bump) or mostly central (left bump). The segregation index is defined as the separation between the clusters (dashed line), divided by the square root of the average of the half widths of the clusters (full widths are solid lines). (**E**) Marginal arbor (green) and central arbor (red) of example GCs (aspect ratio of the cells are distorted for visualization). The IPL depths of the marginal-central boundaries are denoted by *d*_1_ and *d*_2_. (**F**) GC marginal-central segregation index is maximized for *d*_1_ and *d*_2_ at the SAC depths (dashed lines, same values used in **D**). (**G**) Average stratification profiles (linear density of arbor volume vs. IPL depth) of BC types. (**H**) BC types belong to four high-level BC clusters created by inner-outer and marginal-central splits. (**I**) Average stratification profiles (linear density of arbor length vs. IPL depth) of six high-level GC clusters. (**J**) High-level hierarchical clustering of GCs. See also Figure S1.

If we now vary the location of the inner-outer boundary, we obtain a family of histograms (not shown). Each histogram can be summarized by a numerical index of segregation, defined as the separation between cluster centers divided by their average width (Methods, Fig. 3D). The segregation index is graphed versus the location of the inner-outer boundary in Fig. 3C. It turns out that arbor segregation is maximized at 0.47 IPL depth; this was the value used for the histogram in Fig. 3A. The global maximum at 0.47 IPL depth is the On-Off boundary, in the sense that On and Off BC types lie on either side, as will be explained more fully later on. Therefore our arbor segregation principle reproduces the Famiglietti and Kolb [1976] two-way division of the IPL into sublamina *a* and *b*. The novelty is that this division emerges from a purely anatomical principle, without use of physiological information.

Flanking the global maximum at 0.47 IPL depth, there are also local maxima at 0.28 and 0.65 IPL depth (Fig. 3C). The local maximum at 0.28 coincides with the Off SAC depth. The local maximum at 0.65 is very close to the On SAC depth of 0.62 (see Methods about the small deviation). It should be noted that SAC-specific staining has become the most popular landmark for IPL depth [Manookin et al., 2008, Siegert et al., 2009]. The arbor segregation principle supports the idea that SACs are more than merely convenient landmarks; they are objective divisions in the IPL.

To summarize, BC arbor segregation supports the division of the IPL into four sublaminae, which we will call outer marginal, outer central, inner central, and inner marginal. The inner-outer boundary corresponds to the On-Off boundary and the marginal-central boundaries are at the SACs. Further subdivision of the inner marginal sublamina to create a total of five sublaminae is also defensible but less convincing (Methods).

We now examine whether GC arbors segregate across the four sublaminae. It turns out that GC arbors do not segregate well across the inner-outer boundary (Fig. S1B). This is not surprising, given that so many On-Off GC types are known. However, good segregation is obtained if we consider the difference between the amount of arbor in the two marginal sublaminae minus the amount of arbor in the two central sublaminae (Fig. 3D, E). The optimal locations for the marginal-central boundaries are exactly at the On and Off SAC depths (Fig. 3F). For simplicity, the preceding analysis omitted the On-Off and On DS cells [Barlow and Levick, 1965, Sabbah et al., 2017]. The full analysis including the DS cells is slightly more complex while preserving the main findings (Fig. S1D-F and Methods).

## Hierarchical clustering

By dividing the IPL into four sublaminae, the preceding section revealed a difference in BC and GC organization. Marginal-central segregation is stronger for GCs, while inner-outer segregation is stronger for BCs. These splits can be used as starting points for divisive hierarchical clustering of cells. For BCs, the inner-outer split of Fig. 3A is followed by marginal-central splits yielding four high-level clusters (Fig. 3H). Each of these four clusters can be further subdivided into BC types using anatomical criteria described previously [Helmstaedter et al., 2013, Kim et al., 2014, Greene et al., 2016]. The three boundaries between the four sublaminae are visible as “notches” in a graph of the average stratification profiles of the BC types (Fig. 3G). As mentioned earlier, the inner-outer boundary divides the On BC types (BC5-9) from the Off BC types (BC1-4). According to recent data, the On-Off distinction is essentially binary for mouse BCs [Franke et al., 2017].

For GCs, a high-level clustering also follows from the four sublaminae (Fig. 3J, Item S1). We first separate the DS cells based on their strong co-stratification with SACs (Fig. S2B, Item S1, split a-1). The remaining GCs are split into marginal and central clusters as in Fig. 3D (see also Item S1, split a-2). The marginal cluster separates into inner and outer clusters (Fig. S2C, Item S1, split a-4). The central cluster separates into inner, outer, and inner-outer clusters (Fig. S2D, Item S1, split a-3). This procedure yields a total of six high-level clusters (Fig. 3J): DS, inner marginal, outer marginal, inner central, outer central, and inner-outer central. The average stratification profiles of the high-level clusters are shown in Fig. 3I. The inner-outer central cells straddle the inner-outer boundary, and are the main cause of poor inner-outer segregation of GCs, which was noted above.

Each high-level cluster was subdivided using a decision tree (Methods), which yielded a total of 47 low-level clusters. The dendrogram of Fig. S2A shows how the low-level clusters emerge from splits of the high-level clusters (see also Item S1). All decisions in the dendrogram are documented in Item S1. We have compiled a one-page gallery (Fig. S3) that illustrates each cluster with a single example cell, and a multi-page gallery (Item S2) that shows all cells sorted by cluster. All cells and clusters can be viewed interactively at the Eyewire Museum.

Each decision in the tree was made by thresholding an anatomical quantity, which was typically some percentile of the stratification profile (Fig. S4C, D). In some cases we restricted the stratification profile to a particular range of IPL depths and renormalized its area to unity (Fig. S4E, F). This was usually to exclude the dendritic trunks, which appeared to contribute noise to the classification. Finally, some decisions relied on soma size (Fig. S5A-H), SAC contact analysis (Fig. S6), and arbor density and complexity (Fig. S5I-K).

Figure 4A summarizes the anatomy of each cluster by its stratification profile averaged over cells within the cluster. The stratification profiles are grouped by their membership in the six high-level clusters of Fig. 3I. The physiology of each cluster is summarized by its “temporal response function,” defined as visual response versus time, averaged over trials, directions of the moving bar (Fig. 1E), and cells within the cluster. All temporal response functions are normalized in the graphs so that their minimum and maximum values are the same. In reality, response amplitudes varied greatly across clusters, and some clusters responded only very weakly. For example, the 1ws temporal response function is very noisy, because its light-evoked response was so weak, and was averaged over only two cells. More detailed information about visual responses can be found in Item S2 and S3. It should be emphasized that the visual responses were not used at all to define the anatomical clusters. The average visual responses in Fig. 4A were computed after the clustering was complete.

**Figure 4:**
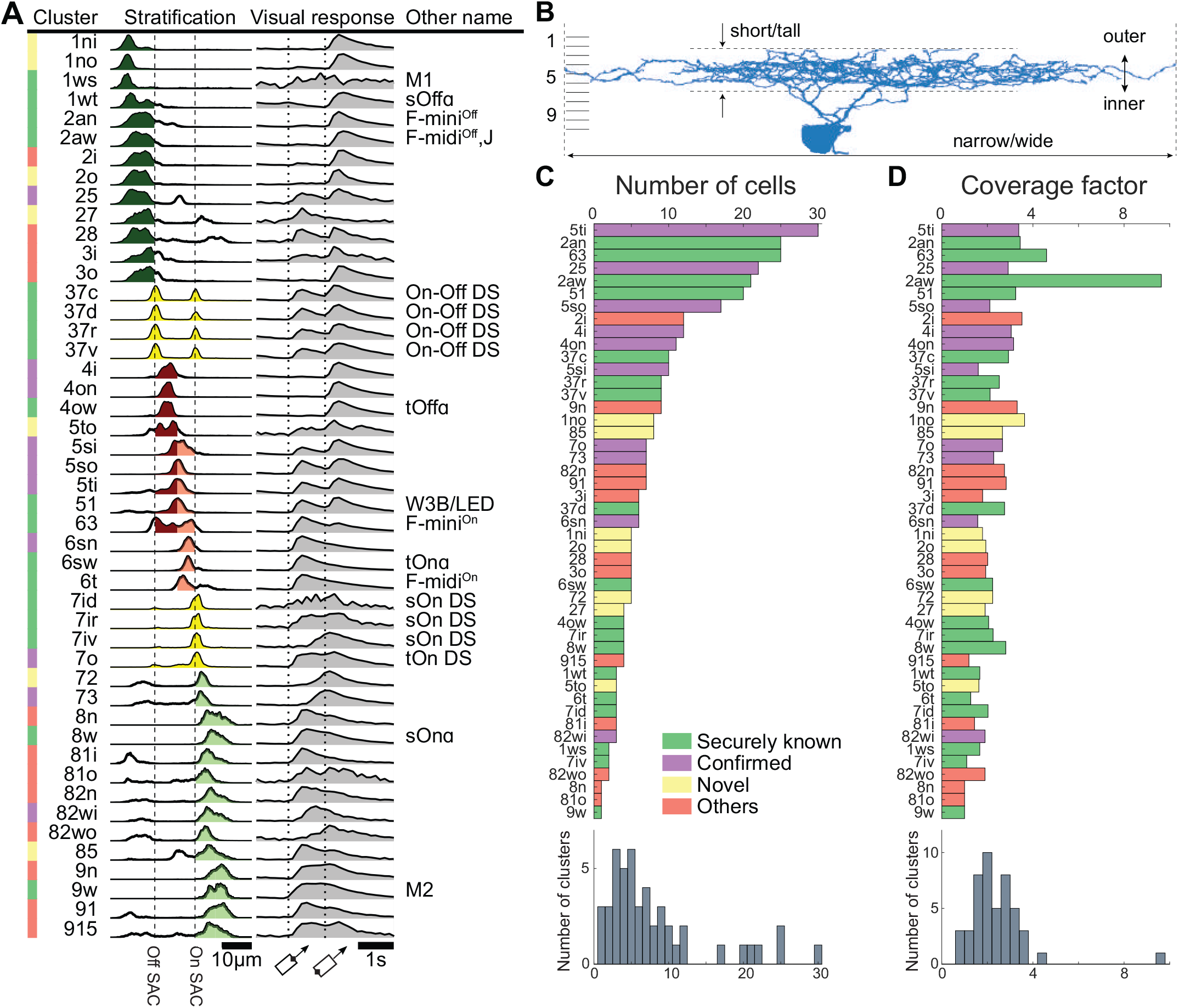
Classification of Ganglion Cells. (**A**) Summary of clusters with anatomical name, stratification profile, and temporal response function defined in Fig. 1E. Alternative names in black are “securely known” types (see main text for definition). (**B**) Each cluster name begins with a number in the range 1-9 indicating which tenth of the IPL depth contains the most stratification profile area. More numbers are appended for multistratified clusters. Letters (s, t, n, w, o, i, a) are added to distinguish between clusters with similar stratification, where “a” denotes asymmetric arbor. (**C**) Number of cells in each cluster. (**D**) Coverage factors. See also Figures S2, S3, S4, S5, and S6.

Our nomenclature for the clusters uses numbers and letters to indicate anatomical properties (Fig. 4B). The name of each cluster begins with one or more integers in the range 1 to 9 that roughly describe the stratification profile. The first number encodes the location of the global maximum of the average stratification profile, when computed over ten bins corresponding to 10 sublayers of the IPL with equal thickness. Added numbers encode the location(s) of a local maxima if they exist. (No maxima were found in the tenth bin.) Letters are added as suffixes to indicate features that distinguish clusters with similar stratification profiles. The division into 10 sublayers is merely a convention, unlike our earlier objective division into four sublaminae.

For any given cluster, one might want a measure of confidence that the cluster is actually a pure GC type. There are several ways to obtain this. First, our clustering procedure is “transparent” in the sense that every decision, while expressed as a computational algorithm, can also be understood and examined by a human. By examining Item S1, one can trace the hierarchical sequence of decisions that lead to any given cluster, which yields some qualitative impression of confidence in the cluster.

Second, many of our clusters can be externally validated because they correspond well with “securely known” types, defined as those that have been extensively characterized by a combination of molecular, physiological, and anatomical techniques [Sanes and Masland, 2015, Rousso et al., 2016]. Correspondences are provided in Fig. 4A (see Methods for detailed justifications). Our 1wt, 4ow, and 8w correspond to the classical alpha types [Pang et al., 2012], and our 6sw corresponds to the nonclassical transient On alpha type [Krieger et al., 2017]. Our 37c, 37d, 37r, and 37v correspond to the On-Off DS types, and 7ir, 7id, and 7iv correspond to the classical On DS types. Our 1ws and 9w correspond to the M1 and M2 melanopsin types. Our 51 corresponds to the W3B type [Zhang et al., 2012]. Our 2an, 63, and 6t correspond to F-mini^Off^, F-mini^On^, and F-midi^On^ of Rousso et al. [2016].

## Density conservation principle

For a third measure of confidence, we would like some quantitative validation procedure that is “internal,” meaning that it depends only on information within the dataset. This could conceivably come from the “mosaic principle,” according to which the cell bodies of a GC type are arranged as if they repel each other [Wassle and Riemann, 1978]. Mosaic analysis utilizes the locations of cell bodies, which today are typically obtained using molecular labeling of a GC type [Kim et al., 2008, Huberman et al., 2008, Zhang et al., 2012, Rousso et al., 2016].

Because our reconstructions have both cell bodies and dendritic arbors, it would be more powerful to use the “tiling principle,” according to which the dendritic arbors of a GC type should “tile” the retina with little overlap, almost like the tiles of a floor. Tiling is traditionally quantified by the coverage factor, which is defined as the average number of arbors that cover a retinal location. Perfect tiling would yield a coverage factor of 1. However, there have been many reports of GC types with coverage factors markedly greater than one [Wassle et al., 1981, Stein et al., 1996]. More recently, genetic techniques have been used to verify that GC types can exhibit high coverage factors while still satisfying the mosaic principle [Zhang et al., 2012, Rousso et al., 2016]. For our GC clusters, the median coverage factor is between 2 and 3 (Fig. 4D), indicating substantial overlap between neighboring arbors.

To illustrate violation of the tiling principle, Fig. 5A shows how the arbors of an example cluster cover the retina. Each arbor is represented by its convex hull, and from the overlap between hulls we can see that the coverage of a retinal location can be as high as 5 (Fig. 5B). However, it is not the case that a region covered by 5 hulls contains 5 × more arbor than a region covered by 1 hull. On the contrary, the aggregate arbor density is almost independent of coverage (Fig. 5C). This example suggests that coverage as quantified by convex hull overlap can be misleading.

**Figure 5:**
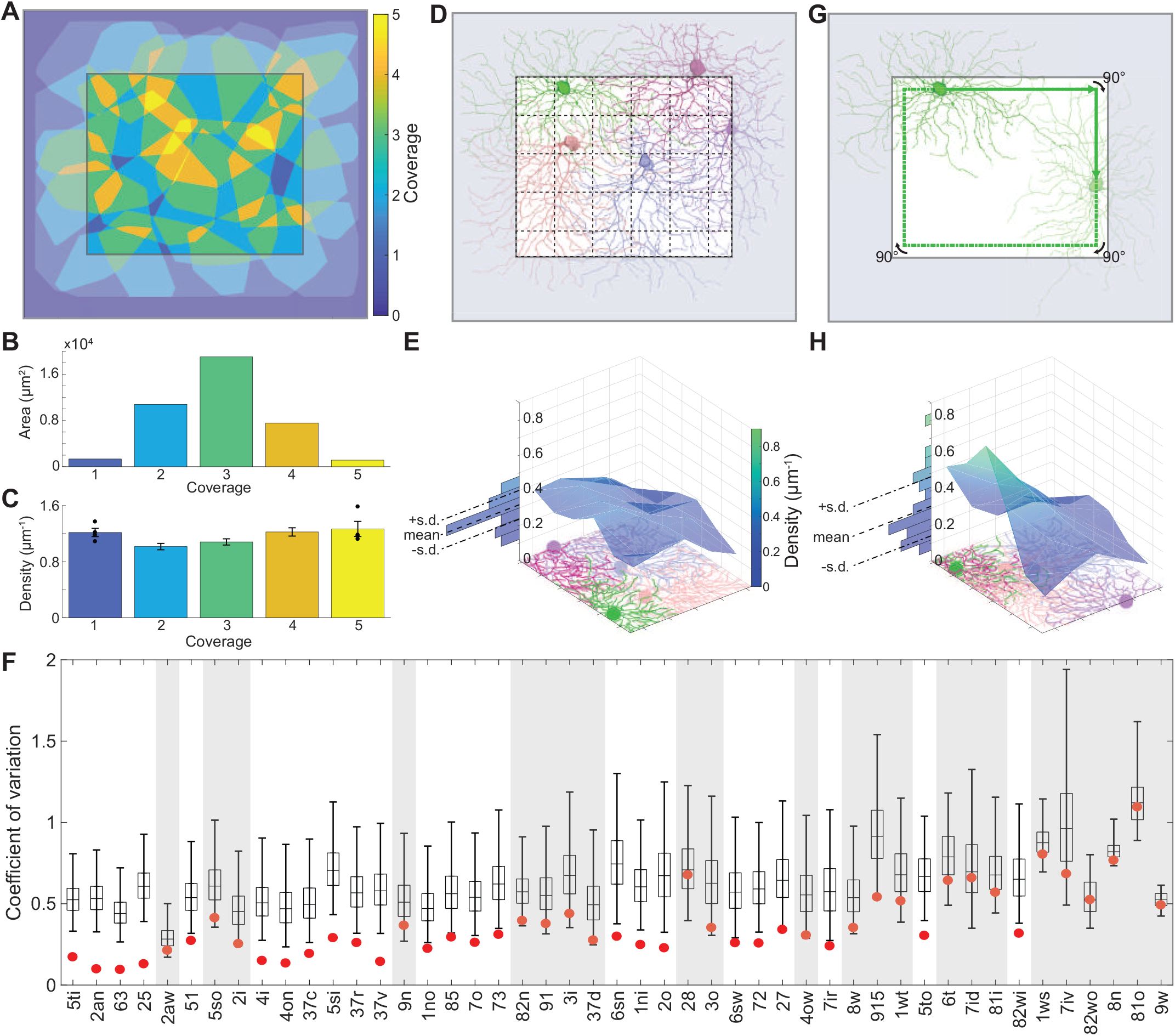
According to Our Density Conservation Principle, the Arbors of a GC Type Should Have an Aggregate Density that is Approximately Uniform. (**A**) Arbor convex hulls of an example cluster (25) overlap substantially. Colors indicate how many hulls cover each retinal location inside the crop region. (**B**) Retinal area versus coverage inside the crop region. Each bar represents the area devoted to the corresponding color/coverage in the crop region. (**C**) The aggregate arbor density of the cluster varies relatively little with coverage. Each bar represents the density within the area devoted to the corresponding color/coverage in the crop region (standard error, *n* = 4, 19, 33, 20, 4). (**D**) The crop region is divided into grid boxes, and the aggregate arbor density is computed for each box, as illustrated for an example cluster (6sw). (**E**) The aggregate arbor density is close to uniform across the crop region, as quantified by the coefficient of variation (standard deviation divided by mean). (**F**) The density conservation test is satisfied by a cluster (non-shaded) when the coefficient of variation is significantly smaller for the real configuration (red dot) than for 99% of all randomized configurations (99/1 percentiles, black bar; quartiles and median, box; *n* = 10,000). (**G**) To test statistical significance, the arbors of a cluster are randomized by relocating the soma somewhere on its “orbit” (green line) and rotating the arbor to have the same orientation relative to the nearest side of the retinal patch. (**H**) The aggregate arbor density typically varies more after randomization. Example cluster is 25 in **A-C** and 6sw in **D**, **E**, **G**, **H**.

Motivated by this example, we propose that the arbors of a type add up to roughly uniform density across the retina. We call this the “density conservation principle,” and it reduces to the traditional tiling principle for the special case of arbors with uniform density within their convex hulls. For arbors that vary in density across their convex hulls, our new principle is compatible with arbor overlap. We have found a prior qualitative report of density conservation in the literature [Dacey, 1989], and related arguments have been made about overlap between GC receptive fields [Borghuis et al., 2008]. Here we present the first quantitative analysis of density conservation, and investigate its universality by applying it to all our GC clusters.

We first defined a central “crop region” in e2198 (Fig. 5D). Cropping excluded the parts of e2198 near the borders, which are expected to have lower aggregate arbor density because we did not reconstruct neurites of cells with their somas outside e2198. The crop region was divided into a grid of boxes (Fig. 5D). In each grid box, we computed the aggregate arbor density. Then we computed the coefficient of variation (standard deviation divided by mean) of the aggregate arbor density across the grid boxes (Fig. 5E). We expected the coefficient of variation to be small, and indeed it was for many cells (Fig. 5F).

To assess statistical significance of a small coefficient of variation, we created an ensemble of randomized configurations from the original cluster. The soma positions and arbor orientations were randomized in a way that left the aggregate arbor density in the crop region roughly constant (Fig. 5G and Methods). The coefficient of variation of the aggregate arbor density was calculated for each randomized configuration (Fig. 5H). We deemed a small coefficient of variation to be statistically significant if it had less than 1% probability of emerging from the randomized ensemble.

We found that 24 of the 47 clusters exhibit statistically significant density conservation (Fig. 5F). We also examined securely known types that failed the density conservation test, and found that these failures were clusters containing relatively few cells (Fig. 4C). For example, 37d is the least numerous of the four On-Off DS types in our sample. It contains few cells (6) and just barely fails the test. 4ow is the second least numerous of the four alpha types in our sample. It also contains few cells (4) and just barely fails the test. Both types might have passed the test had our sample been larger. So there is no strong violation of the conjecture that density conservation is universal for all GC types.

Even if a necessary criterion, density conservation cannot by itself be a sufficient criterion for a pure GC type. To see why, consider a thought experiment in which two pure types exactly satisfy density conservation. A mixture of the two types will also exactly satisfy density conservation. However, we should be able to detect a mixture by its abnormally large coverage factor, which will be the sum of the coverage factors of the pure types.

Therefore, we propose that a cluster is internally validated as a type if (1) it satisfies density conservation and (2) its coverage factor is in the normal range. This range is roughly 1.5 to 3.5 for our clusters (Fig. 4D), which is consistent with many reports of GC coverage factors in the literature. (The coverage factors lower than 1.5 in Fig. 4D are likely underestimates, because they are mainly for clusters containing few large cells, for which the coverage factor computation is corrupted by edge effects.)

If we take the union of our internally validated types with the securely known types, we end up with a total of 33 clusters that are validated as types, plus one cluster (2aw) that appears to contain two securely known types (F-midi^Off^ and J). These contain 84% of all cells in the sample of 381 (Fig. 4C). Further discussion of the 2aw and 63 clusters can be found in the Methods.

## Novel ganglion cell types

While the present work focuses on general principles of IPL organization, as a bonus it also yields six particular GC types that appear to be novel. For six of our internally validated clusters (1ni, 1no, 2o, 85, 27, 5to in Fig. 6), we have been unable to find unambiguous correspondences with previously published types. Outer marginal types 1ni and 1no look remarkably similar in Figs. 6A and 6B, and are novel types that co-stratify with 1ws (M1 melanopsin). Our clustering procedure separates 1ni and 1no based on a small but systematic difference in their stratification profiles (Fig. 6C and Item S1, split e-7). Their average temporal response functions also differ slightly (Fig. 6C).

**Figure 6:**
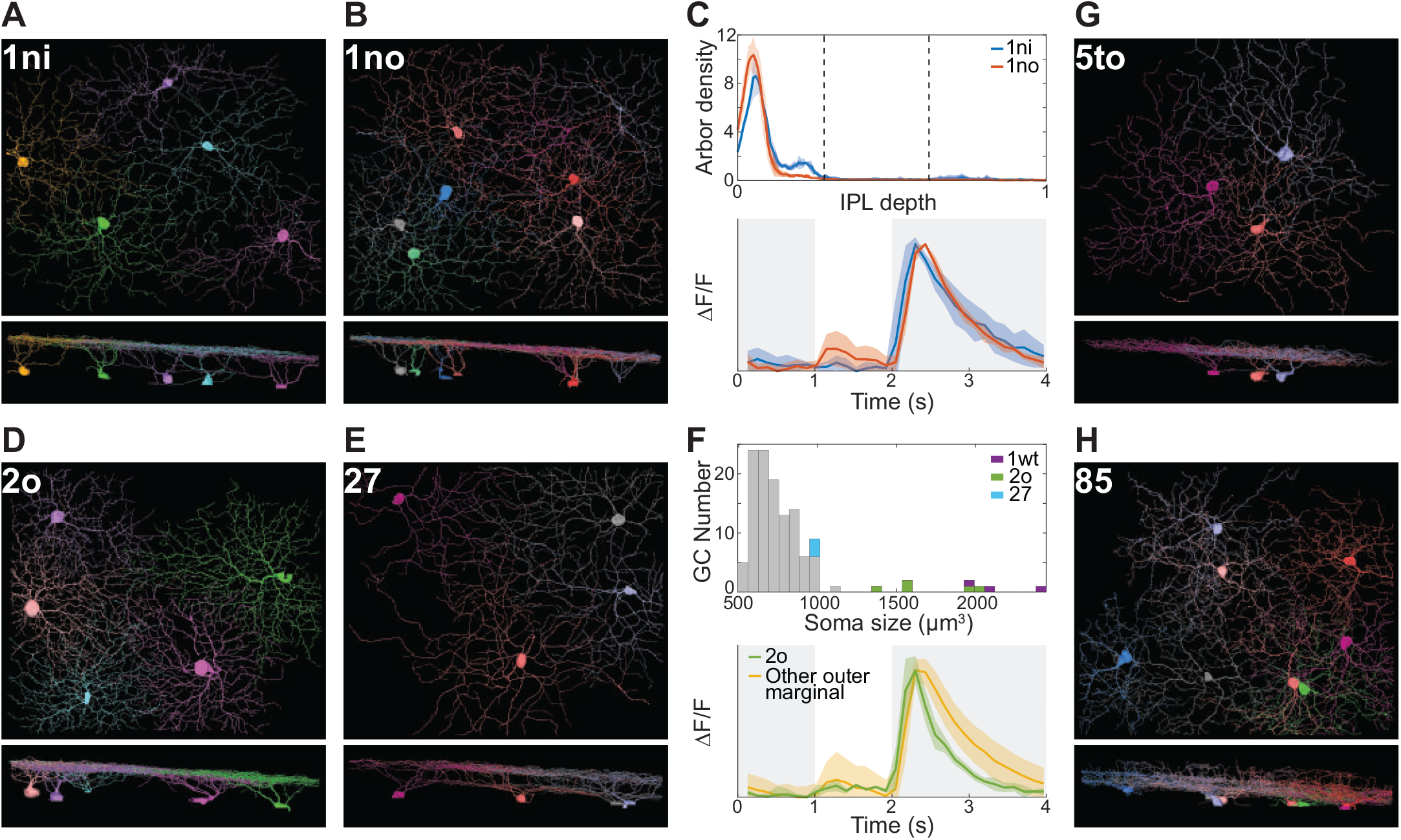
Novel Ganglion Cell Types, Views Along the Light Axis and a Tangential Axis. (**A-C**) 1ni and 1no are types with very similar stratification profiles (**C**, top) and temporal response functions (**C**, bottom). (**D-F**) 2o and 27 are outer marginal types. Histogram of soma size for outer marginal cells shows that 2o somas are much larger than those of 27 and other typical cells, and smaller than 1wt (transient Off alpha) somas (**F**, top). The Off response of 2o decays more rapidly than that of other outer marginal cells (**F**, bottom). (**G**) 5to looks monostratified in the tangential view but its stratification profile (Fig. 4A) is surprisingly complex. (**H**) 85 stratifies throughout the inner IPL but also extends sparse branches towards the INL. Shaded regions around curves in **C** and **F** represent standard deviations.

Outer marginal types 2o and 27 are depicted in Figs. 6D and 6E. The 2o soma is almost as large as that of the classical alpha types, which have the largest somas of all GCs (Fig. 6F). The 27 soma is of more typical size (Fig. 6F). Of the outer marginal cells, 2o exhibits the most transient Off response (Fig. 6F).

Outer central type 5to appears monostratified in the tangential view of Fig. 6G. However, its average stratification profile is relatively broad and contains more than one peak (Fig. 4A). Multiple peaks can also be seen in the stratification profiles of individual 5to cells (Item S2 and Museum). Inner marginal type 85 arborizes throughout the inner IPL, but also extends sparse branches towards the INL (Fig. 6H). Its average stratification profile (Fig. 4A) exhibits three peaks in the inner marginal, inner central, and outer marginal sublamina. Both 5to and 85 show that the stratification profile can be very rich in information.

These and all other clusters can be interactively viewed in the Eyewire Museum, which provides search bar access to reconstructions (Fig. 2B), stratification profiles (Fig. 2F), and visual responses (Fig. 2G, H).

Nine of our internally validated types (5ti, 5so, 5si, 4i, 4on, 6sn, 7o, 73, 82wi) correspond with GC classes that were recently defined by physiological and anatomical techniques [Baden et al., 2016, Jacoby and Schwartz, 2017, Mani and Schwartz, 2017, Sabbah et al., 2017] but have not yet been confirmed as pure types by molecular techniques and mosaic analysis. Our density conservation test (Fig. 5F) provides supporting evidence that these classes are indeed pure types. Our 5ti, 5so, and 5si co-stratify with 51 (W3b), and correspond to the HD family of Jacoby and Schwartz [2017]. Our 4i and 4on co-stratify with 4ow (transient Off alpha), and our 6sn co-stratifies with 6sw (transient On alpha). These may correspond with “mini alpha” types identified by Baden et al., 2016. Our 7o corresponds to the nonclassical transient On DS cell found by BadenBaden et al., 2016Baden et al., 2016 and Sabbah et al. [2017]. Our 73 corresponds to the On delayed cell as defined by Mani and Schwartz [2017]. Our 82wi corresponds to the vertically orientation selective cell studied by Nath and Schwartz [2016]. Some of the confirmed types can also be found in the survey of Helmstaedter et al. [2013]. One of our internally validated types (25) confirms a type in Helmstaedter et al. [2013] that has not yet been identified by physiologists. 25 is the fourth most numerous cluster in our sample (Fig. 4C). More detailed evidence for the above correspondences is given in the Methods.

## Sustained vs. transient

Our distinction between marginal and central GC clusters occurs at the top of the GC hierarchy (Fig. 3J and Fig. S2A), so it seems fundamental. Could this anatomical distinction have functional significance? Physiologists classify retinal neurons as sustained or transient, mainly based on duration of response to a sudden change in illumination [Cleland et al., 1971]. A previous study combined electrophysiology and light microscopic anatomy to provide evidence that central GCs are transient, while marginal GCs are sustained [Roska and Werblin, 2001]. Our dense sample of GCs provides an opportunity to systematically characterize differences in sustainedness.

For each high-level cluster we averaged the temporal response functions of all cells in the cluster. The average response of the inner marginal cluster is markedly more sustained than that of the other high-level clusters (Fig. 7A). We then quantified the sustainedness of each cell by the value of its temporal response function at a fixed time after nominal stimulus onset, relative to the peak value (Fig. 7B and Methods). Inner marginal cells are significantly more sustained on average than the cells in the other high-level clusters (Fig. 7C). Differences between other clusters are not statistically significant.

**Figure 7:**
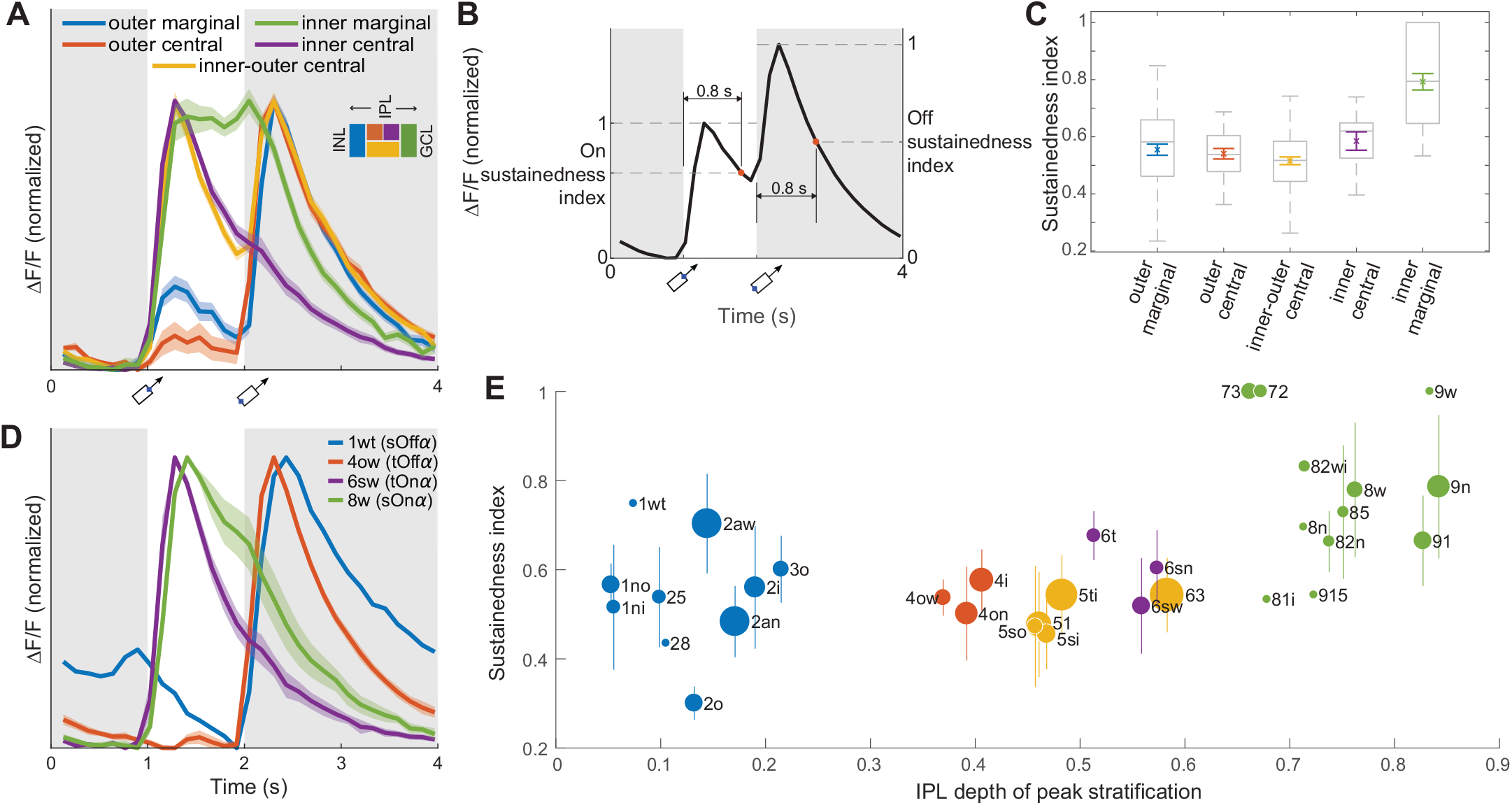
Sustainedness of Visual Responses and Dendritic Stratification. (**A**) Average temporal response function (Fig. 1E) for high-level GC clusters (Fig. 3I). Each response function is averaged over cells in a cluster, and normalized to have the same maximum and minimum. The inner marginal cluster is markedly more sustained than the others. Shading indicates standard error (*n* = 102, 26, 78, 12, 55 for outer marginal, outer central, inner-outer central, inner central, inner marginal). (**B**) The sustainedness index is defined as the response at 0.8 sec after nominal stimulus onset, divided by peak response in the 0.8 sec interval. (**C**) The cells in the inner marginal cluster are significantly more sustained than cells in the other clusters (ANOVA with post hoc *p <* 0.01). The differences between other clusters are not statistically significant. Colored, mean and standard error; grey box and bars, quartiles, median, and extrema. (**D**) Marginal is more sustained than central for the four alpha types. Shading indicates standard error of the mean (*n* = 4, 5, 4 for 4ow, 6sw, 8w). There is no standard error for 1wt, because only a single 1wt cell had calcium signals. Applying *t*-tests to sustainedness indices yield *p* = 0.02 for 1wt against 4ow, and *p* = 0.03 for 8w against 6sw. (**E**) Sustainedness index for cells with high response quality (Methods, Eq. 4), sorted by clusters. Bars indicate standard deviations for the clusters, except for clusters containing only a single cell with high response quality (1wt, 28, 81i, 8n, 915, 9w). Dot area indicates the number of cells in the cluster; the largest dot (63) represents 18 cells. See Figure S7 for size-adjusted sustainedness index for individual cells.

One might worry that the temporal resolution of calcium imaging is inadequate for revealing sustained-transient differences. The four alpha types are a counterexample to this idea, because they exhibit sustained-transient differences (Fig. 7D) consistent with those that have been observed using electrophysiology [Pang et al., 2012, Krieger et al., 2017].

We refine the analysis by plotting the average sustainedness for the cells of each low-level cluster versus its peak stratification (Fig. 7E). For comparison, we also plot the average temporal response function for each low-level cluster (Fig. S7A). The numbers for any individual cluster should be taken with caution, especially as some clusters have not been validated as types. The sustainedness of the outer marginal cells spans a wide range; 1wt is most sustained, while 2o is most transient. The sustainedness of the inner marginal cells also spans a wide range. Given this diversity, the idea that marginal cells are more sustained than central cells may be simplistic.

One concern about the analysis is that our stimulus is a moving bar, while a flashed stimulus is customarily used for the sustained-transient distinction. For a moving stimulus, the trailing edge leaves the receptive field later for an arbor with larger diameter. This might cause large cells to spuriously seem more sustained than they actually are. To control for this possibility, we recalculated the sustainedness indices after translating the response of each cell backwards in time by an amount equal to its cluster-specific arbor radius divided by the stimulus speed (Methods). The result is shown in Fig. S7B, which looks very similar to Fig. 7E.

## Discussion

In light of our findings for GCs, it is helpful to reexamine the analogous claim that marginal BCs are sustained while central BCs are transient [Borghuis et al., 2013, Baden et al., 2013]. For a full field flashed stimulus, inner marginal BCs are the most sustained cluster; the differences between the other three clusters are more minor (Extended Data Fig. 6m of Franke et al., 2017). This finding is strikingly analogous to our own finding for GCs, and would be predicted from the hypothesis that GCs inherit their sustainedness from their BC inputs [Awatramani and Slaughter, 2000].

The conventional wisdom that marginal BCs are more sustained than central BCs still appears valid, if the comparison is restricted to inner cells only or outer cells only [Franke et al., 2017]. But it would be misleading to say that marginal is unconditionally more sustained than central, as inner central BCs can be more sustained than outer marginal BCs [Franke et al., 2017], and the same may be true for our GCs (Fig. 7E). One caveat is that sustained versus transient could depend on the specifics of the stimulus. For example, Franke et al., [2017] find that inner central cells are more transient with a full field than a local stimulus. A second caveat is that there may be heterogeneity across the types within a high-level cluster. For example, BC9 is markedly more transient than other inner marginal BC types for a full field stimulus [Franke et al., 2017]. Heterogeneity is likely even greater for GCs, which come in many more types than BCs. A third caveat is that sustainedness may depend on whether conditions are photopic or scotopic [Grimes et al., 2014]. All of these caveats support the overall conclusion that sustained versus transient is more complex and subtle than a simple dichotomy.

Our purely anatomical subdivision of the IPL into marginal and central sublamina supports the idea that using SACs as landmarks is fundamental, and not merely a convenience made popular by the ease of ChAT staining. It has been proposed that SAC arbors could serve as a scaffold for development of some types of GC arbors [Stacy and Wong, 2003], so the marginal-central division is potentially relevant for neural development.

We have proposed a density conservation principle to replace the tiling principle for GC types. Density conservation makes functional sense as it would serve to make uniform the density of synapses from each BC type to each GC type. The tiling principle can be fulfilled if dendrites of different cells repel each other during development [Grueber and Sagasti, 2010]. We speculate that density conservation could be established by a developmental process in which homotypic dendrites repel each other without regard to whether they belong to the same cell or different cells.

Of the 47 clusters that we identified, 24 were internally validated by the density conservation test (Fig. 5F). Most of the remaining clusters contain too few cells to satisfy the criteria with statistical significance, but some of them can be externally validated because their properties match “securely known” types that have been published previously. If we take the union of internally validated types and securely known types, we end up with a total of 35 types. This lower bound on the number of GC types is consistent with the recent identification of 35 to 50 clusters in the large-scale physiological survey of mouse retinal GCs mentioned previously [Baden et al., 2016].

## Author Contributions

KLB acquired the e2198 dataset, and performed preliminary analysis of the calcium imaging data (including manual ROI detection). JAB analyzed arbor segregation and density conservation with input from SM, JSK, and NLT. SM analyzed sustained vs. transient responses. Eyewirers reconstructed neurons with supervision from RP, MS, CD, DLJ, and DB. RP devised strategies for expert correction of crowd wisdom (“reaping”). ALRS managed Eyewire operations and analyzed efficacy of crowdsourcing techniques. SM created code for detecting reconstruction errors by finding collisions between neurons. JSK curated neuron reconstructions, and performed the computational flattening. JSK and SM found correspondences between calcium ROIs and EM cell bodies. JAB, SM, and JSK performed hierarchical clustering. JAB and NLT transformed 3D reconstructions into skeletons. SM performed SAC contact analysis. NLT segmented cell bodies with help from M. Moore, and quantified arbor asymmetry. JSK computed convex hulls. IT, WMS, and ADN created the online museum. WMS created the system by which scythes mark branches as complete, with help from K. Radul. WMS added support for multiple languages with help from M. Balkam and K. Radul. NK devised a new point system for gameplay that better incentivized accuracy, and implemented a chatbot that helped with community management. CSJ added Eyewire features used for collaboration with KT. JMP worked on KT promotions with help from H.-J. Park, S.-H. Seo, J. Hong, E. Bae, and S.-B. Yang. HSS wrote the paper with help from JAB, SM, JSK, NLT, NK, WMS, ALRS, JMP, and KLB.

## Acknowledgments

We are grateful to W. Denk for providing the e2198 dataset. We thank Dr. Chang-kyu Hwang for originating the idea of collaboration between KT Corporation and Eyewire on “citizen neuroscience,” and H. Park for helping to make the connection. S. Caddick and D. Feshbach provided guidance to WiredDifferently. We thank S. Ströh for help in identifying correspondences with cell types in the literature. We are grateful to M. J. Greene for helping start the contact analysis and skeletonization. We thank M. Balkam and K. Radul for assistance with Eyewire in the early stages of this investigation. M. Kim, K. Lee, and D. Ih helped manage the Korean Eyewire community. B. Paiva contributed Eyewire logo animations. C. Xiang and N. Benson created badges and art for promoting competitions. N. Friedman created educational material for the Eyewire blog and wiki. C. O’Toole and D. Sparer promoted Eyewire on social media and blogs. M. Akasako contributed material on cell types to the wiki. We benefited from interactions with P. Berens, M. Berry, D. Berson, B. Borghuis, T. Euler, J. Homann, M. Meister, T. Schmidt, G. Schwartz, and J. Sanes. Research was supported by the Gatsby Charitable Foundation, NINDS/NIH (U01NS090562 and 5R01NS076467), DARPA (HR0011-14-2-0004), ARO (W911NF-12-1-0594), IARPA (D16PC00005), KT Corporation, and the Amazon Web Services Research Grants Program. JSK acknowledges support from the Korea Brain Research Institute basic research program funded by the Ministry of Science, ICT, and Future Planning through award 2231-415.

## STAR Methods

### Countdown to Neuropia

On August 12, 2014, KT Corporation and Eyewire signed a memorandum of understanding in which KT pledged to “fulfill its corporate social responsibility by mobilizing Korean people to participate in Eyewire, thereby using telecommunications technology to advance neuroscience research for the benefit of all humanity.” The signing ceremony was held at KT Olleh Square in central Seoul and was covered by over 40 mass media outlets including television, newspapers, and websites. The memorandum laid out the KT-Eyewire plan for Countdown to Neuropia, a campaign to reconstruct ganglion cells in the e2198 dataset.

To prepare for the Countdown, the Eyewire site was translated into Korean and a separate Korean chat channel was created and moderated by Korean-speaking lab members. The Countdown officially launched on October 10, 2014 with four months of nationwide television advertising. Banner ads were posted on the main pages of various portal websites. KT created a microsite to promote Eyewire with prizes. More than 2,600 players participated in a six-week long competition which awarded monetary prizes totaling $50,000. A few top players won the opportunity to visit Eyewire headquarters and several U.S. universities. In addition, KT mobilized an existing group of 280 students from 40 colleges who serve as brand ambassadors of KT. From October to December 2014, these students (known as “Mobile Futurists”) both played Eyewire to win prizes, and publicized Eyewire at their colleges. From March to July 2015, competitions were organized at five high schools. A total of 511 students competed to win weekly and overall best rank in their respective schools. KT promotions closed in July 2015 with the completion of Phase 3 of the Countdown.

During the KT promotions from October 2014 to July 2015, there were an estimated 4,271 Korean and 9,532 non-Korean participants. (Participant is defined as a player who submitted a non-tutorial cube. Korean vs. non-Korean was inferred based on IP address, language setting of web browser, and participant lists of KT promotions.) 13,878 Korean players registered, 38 times the number from October 2013 to July 2014. Korean players completed 879,713 cubes, 33% of the total cubes played during the period.

The e2198 GCL patch was divided into four zones with borders defined by three concentric squares with side lengths of 0.05, 0.1, and 0.2 mm. There were 27, 79, 242, and 456 cell bodies in Zones 1 through 4, enumerated from inside to outside. In the first three zones, 7, 16, and 62 had already been reconstructed for previous studies [Kim et al., 2014, Greene et al., 2016] and other preliminary studies in lab. The remaining 20, 63, and 180 cells were reconstructed in three successive Countdown Phases.

The Countdown started in October 2014, and Phases 1 through 3 concluded on November 2014, Feburary 2015, and July 2015. The end of Phase 3 was celebrated by the release of a video showing all cells in Zones 1 through 3 (Movie S1). After cells were reconstructed, GCs were distinguished from amacrine cells (ACs) by the presence of axons. The total numbers of GCs in the first three zones turned out to be 13, 43, and 112 (Item S2).

Since Zone 4 contained so many cell bodies, we decided to restrict reconstructions to GCs only. We inspected 456 candidate cell bodies in Zone 4, and identified 228 as GCs by detecting axons. Of these 228 cells, 60 had already been reconstructed previously. The remaining 168 “bonus cells” were reconstructed in the fourth and last “Bonus Phase” of the Countdown, which concluded in November 2015.

GC axons can be challenging to detect when they branch from dendrites rather than directly from the soma. Searching for axons in Zones 1 to 3 was done after all cells were fully reconstructed, and was therefore likely more reliable than identification of GCs in Zone 4, which was done prior to reconstruction. Therefore the false negative rate for Zone 4 GCs may be higher than in the first three zones.

Overall, 396 GCs were reconstructed, with a total path length of roughly 1.52 m. We excluded 11 cells that were severely cut off by the borders of the EM volume, and four more “weirdos” that may have resulted from developmental abnormalities, leaving 381 GCs for subsequent analysis.

### Computational flattening and downsampling

The IPL in our volume had some curvature and variations in thickness. We computationally flattened the IPL by deforming it so that the Off and On SAC layers became parallel planes [Sümbül et al., 2014, Greene et al., 2016]. Such flattening has previously been shown to increase the reproducibility of stratification profiles [Manookin et al., 2008, Sümbül et al., 2014, Greene et al., 2016]. We adapted the code developed by Sümbül et al. [2014]. Each SAC layer was quasi-conformally mapped to a plane located at its median IPL depth (Fig. S4A). The mappings were extended from the SAC planes to the rest of the IPL by using local polynomial approximations. The transformation was applied to the entire e2198 volume, along with 4× downsampling in each direction.

### Stratification profiles

For BCs, we defined the stratification profile as the linear density of arbor volume as a function of IPL depth [Kim et al., 2014, Greene et al., 2016]. For GCs, we defined the stratification profile as the linear density of arbor length as a function of IPL depth. The different definitions (volume vs. length) were chosen to lessen the contribution of the arbor trunk to the stratification profile. For BCs, the caliber of the trunk is often less than the caliber of the branches, so using volume tends to weight the trunk less. For GCs, the caliber of the trunk is generally greater than the caliber of the branches, so using volume tends to weight the trunk more.

The 3D reconstruction of each GC arbor was automatically transformed into a 1D skeleton (Fig. S4B), and for each skeleton we computed the density of voxels as a function of IPL depth. This “stratification profile” was treated like a probability distribution, with its area normalized to unity.

### Automated skeletonization

GC skeletons were computed from 3D reconstructions as follows. We define two graphs on the voxels of the cell, with edges determined by 26-connectivity. In the undirected graph, the weight of the edge between voxels *ν* and *ν′* is given by the Euclidean distance *d*(*ν*, *ν′*), taking on values 1, 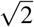, or 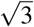. In the directed graph, the weight of the edge from voxel *ν* to *ν*′ is

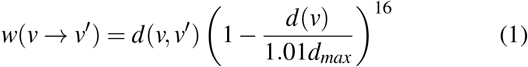

Here *d(ν)* is the Euclidean distance from voxel *ν* to the boundary of the cell, which is known as the “distance boundary field” and computed using a Euclidean Distance Transform algorithm from Maurer et al. [2003]. This procedure ignores voxel anisotropy. The maximum of *d(ν)* over all voxels in the cell is *d_max_*. Equation (1) differs from Sato et al. [2000], who used a sum rather than a product.

A root is selected as the first voxel in the dictionary ordering of the *xyz* voxel locations, which is typically at the end of a dendrite. A destination is chosen as the farthest voxel in the undirected graph. Then the shortest path between root and destination is computed in the directed graph to yield part of the skeleton. Then the undirected graph is modified by removing all points in cubes centered on the skeleton voxels, where the cube at voxel *ν* has length 6*d*(*ν*) + 6. The dependence on *d(ν)* means that the cube is bigger where the dendrite is thicker. The numerical values set the size of “spines” that will be ignored. Based on this modified undirected graph, a new destination point is selected as the farthest point from the root. Then we compute the shortest path from the root to the destination in the directed graph. This process is iterated until no points remain in the undirected graph.

A cell may consist of multiple connected components due to small inaccuracies of the reconstruction process. Each connected component is skeletonized separately, and the stratification profile is computed from the set of skeletons.

### Arbor segregation principle

The optimal location for the outer marginal-central boundary is at IPL depth 0.62 (Fig. 3C), which is the same as the Off SAC depth. The optimal location for the inner marginal-central boundary is at IPL depth 0.65 (Fig. 3C), which is slightly displaced relative to the On SAC depth of 0.62. One possible reason for the asymmetry is the On DS types, some of which have peak stratification close to 0.65. If central BCs synapse onto On DS cells, they would need to extend beyond the On SACs, causing the marginal-central boundary to shift.

We divided the IPL into four sublamina by placing borders located at the three peaks in Fig. 3C. The inner marginal sublamina is larger than the others, so it is natural to ask whether it can be subdivided to create a total of five sublamina as in Cajal [1893]. There is a tiny local maximum at 0.71 IPL depth in Fig. 2c, so one could place a border there to divide BC7 from BC6, BC8/9, and RBC. The segregation is not as good, however, so we prefer the system of four sublamina.

The analysis of GC arbor segregation in the main text omitted the On-Off and On DS cells. If they are included, then Fig. 3F changes to Fig. S1D, which contains two local optima of almost identical quality. One optimum assigns the DS cells to the marginal cluster (Fig. S1E), while the other assigns the DS cells to the central cluster (Fig. S1F). It makes intuitive sense that the DS cells are a borderline case, because they co-stratify with SACs, which are the marginal-central boundaries. Since it is arbitrary whether the DS cells are assigned to the central or marginal cluster, it also seems reasonable to assign them to neither, which was the strategy taken by the simplified analysis in the main text.

### *k*-means clustering in 1D

In our hierarchical clustering, every split was made by applying *k*-means clustering in 1D. We used *k* = 2 for all splits except one, for which we used *k* = 3. The centroids of the clusters were randomly initialized using the method of Arthur and Vassilvitskii [2007].

For some of the high-level splits, we defined the segregation index

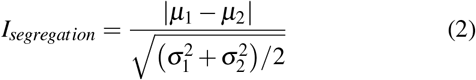

where *μ*_1_ and *μ*_2_ are the centroid locations and 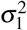 and 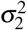 are the cluster variances. The segregation index was averaged over 1000 random initializations of the *k*-means algorithm.

### Hierarchical clustering of GCs

The hierarchical clustering of GCs is depicted by the dendrogram of Fig. S2A. The top levels of the hierarchy are the same as in the smaller dendrogram of Fig. 3J. In the first division, DS cells are separated by cosine similarity with On and Off SACs (Fig. S2B). The remaining cells are separated into marginal and central clusters (Fig. 3D). The marginal cluster separates into inner and outer clusters (Fig. S2C). The central cluster separates into inner, inner-outer, and outer clusters (Fig. S2D). The preceding yields six high-level clusters, which are further divided into 47 clusters based mainly on features computed from the stratification profile (Fig. S4C-F). Soma size, SAC contact, and arbor density and complexity are used for a few divisions.

From the outer central cluster, 5to separates by 10th percentile of IPL depth (split b-1). Then, 4ow separates by its large soma size (split b-2). The remaining cells separate into 4on and 4i based on the difference between the 70th and 15th percentile IPL depths (split b-3).

From the inner-outer central cluster, 63 separates via cosine similarity with the Off SAC stratification profile (split c-1). From the remaining cells, 5si and 5so separate via 5th percentile of IPL depth (split c-2), and are distinguished from each other based on the 80th percentile IPL depth (split c-4). The remaining cells divide into 51 and 5ti by arbor complexity (split c-3).

From the inner central cluster, 6t separates via 95th percentile IPL depth (split d-1). The remaining cells divide into 6sw and 6sn based on soma size (split d-2).

From the outer marginal cluster, 27 and 28 separate via 85th percentile IPL depth of the stratification profile restricted to the marginal IPL (split e-1), and are distinguished from each other via 95th percentile IPL depth (split e-12). From the remaining cells, 1wt and 2o separate based on soma size (split e-2), and are distinguished from each other via arbor complexity (split e-11). 1ws, 1no, and 1ni can be separated via 50th percentile IPL depth (split e-3) because their dendrites are adjacent to the INL. 1ws can be distinguished from 1no and 1ni by arbor density (split e-5), and 1no and 1ni can be separated via 85th percentile IPL depth (split e-7). From the remaining cells, 2an and 25 separate based on arbor complexity (split e-4), and are distinguished from each other by the difference of the 90th and 45th percentile IPL depths (split e-10). From the remaining cells, 3i and 3o separate via the difference between the 80th and 10th percentile IPL depths of the stratification profile restricted to the outer IPL (split e-6), and are distinguished from each other by the difference between the 90th and 10th percentile IPL depths of the same restricted stratification profile (split e-9). The remaining cells divide into 2aw and 2i based on arbor density (split e-8).

From the inner marginal cluster, 8w separates via soma size (split f-1). From the remaining cells, 72, 73, 81o, and 82wo separate by 95th percentile IPL depth (split f-2), and 81o and 82wo separate from 72 and 73 by 80th percentile IPL depth (split f-6). 72 and 73 are distinguished from each other via stratication profile area of central region (split f-9), while 81o and 82wo are distinguished from each other via 25th percentile IPL depth (split f-10). From the remaining cells, 85 separates via the inner central stratification profile area (split f-3). From the remaining cells, 8n,9n, and 9w separate by 5th percentile IPL depth (split f-4), and 8n is distinguished from 9n and 9w by 50th percentile IPL depth in detailed resolution (split f-8). Then, 9n and 9w are distinguished from each other by dendritic field area (split f-13). Among the remaining cells, 915 and 82n have extra arbors in the central region so they are separated via the central stratification profile area (split f-5), and distinguished from each other by the difference between marginal and central stratification profile areas (split f-12). 91 is separated from the rest of the cells by the 50th percentile IPL depth of the stratification profile restricted to the inner region (split f-7). The remaining cells are clustered into 81i and 82wi via 25th percentile IPL depth (split f-11).

### Soma size

Semiautomated reconstruction in Eyewire is based on a convolutional network described previously [Kim et al., 2014]. This convolutional network worked well in the IPL, but was inaccurate in the GCL because of large gaps in boundaries between cell bodies caused by incomplete staining. To segment the GCL, we started with a 2× downsampled image. Then we applied minimum filtering with a 9 × 9 × 1 sliding window and a stride of 5 in all dimensions. This yielded a new image with 10× downsampling in each dimension relative to the original image. The convolutional network was applied, and its output was segmented using a modified watershed transform (http://github.com/seung-lab/watershed and Zlateski and Seung, 2015). A human annotator examined all ganglion cell bodies and removed all but small parts of the primary dendrites. Often no action was required, as almost all of the dendritic arbor of a cell was generally split off from the cell body due to the downsampling.

Our clustering procedure relied on a few other anatomical properties beyond stratification profile. Soma size is easy to quantify, and has long been studied by anatomists. 8w, 4ow, and 1wt are the types with the largest median soma size (Item S4A), and are clear outliers in the distribution of soma sizes (Fig. S5A, B). They correspond to three types of alpha cell that have been identified in mouse based on soma size, branching pattern, and arbor diameter [Pang et al., 2012]. 8w, 4ow, and 1wt also have the thickest primary dendrites (Fig. S5C). 4ow (transient Off alpha) and 6sw (transient On alpha) are just inside the Off and On SAC layers, respectively. 1wt (sustained Off alpha) and 8w (sustained On alpha) are close to the INL and GCL, respectively. The sustained alpha cells are “taller” than the transient alpha cells (see Fig. 4B for definition of “tall”). 1wt has a secondary peak in the stratification profile closer to the Off SAC layer.

After the three classical alpha types, the largest somas are mostly DS cells (Item S4A). A notable exception is 6sw. It has a smaller soma than 4ow, but its arbor is very similar when viewed along the light axis. 6sw corresponds to a fourth alpha type that has recently been genetically identified [Krieger et al., 2017], and to “ON transient, large” G_19_ of Baden et al., 2016. 6sw was presumably not regarded as an alpha cell by Pang et al. [2003] because its soma and primary dendrites, though large, are not as large as those of the classical alpha types.

When 6sw is included, each of the four IPL sublaminae contains one alpha type (Fig. S5C): inner marginal (8w), inner central (6sw), outer central (4ow), and outer marginal (1wt). Within each type, the stratification profiles are highly reproducible (Fig. S5C). When viewed along the light axis, the four types appear very similar to each other (Fig. S5D-G). Each of the types contains 3 to 5 cells that cover the retinal patch (Fig. S5D-G).

2o is another cell type with a large soma, roughly in the same range as 6sw and the DS cells (Item S4A). Soma size is used by our clustering to separate 2o from three other types (2i, 2an, and 2aw) with almost identical stratification profiles.

A number of types are nearly identical to alpha cells in stratification and visual response (Fig. 4A), but have reduced soma size (Item S4A). 4on and 4i are similar to 4ow, 6sn matches 6sw, and 8n matches 8w. These types may correspond to the “mini” alpha cells recently identified by Baden et al., 2016, which have the same visual responses as alpha types but smaller somas and no SMI-32 staining.

### SAC contact analysis

On DS and On-Off DS types separate from each other via SAC contact analysis. The flattened and downsampled EM volume between IPL depths 0.1 and 0.8 was divided into a regular grid of rectangular cuboids. Each cuboid was 15 × 11 voxels (roughly 1 μm) in the tangential plane, and 181 voxels (12 μm) along the light axis. The grid was exactly two cuboids deep along the light axis. One cuboid was outer IPL (depth 0.1 to 0.45) and the other cuboid was inner IPL (depth 0.45 to 0.8).

For each GC, we examined all reconstructed SACs and recorded two sets of SAC voxels. Firstly, we found all *contacting* voxels, defined as SAC surface voxels contacting the GC. Secondly, we found all *collocating* voxels, defined as SAC surface voxels in the grid cuboids occupied by the GC. The contacting voxels are a fraction of the collocating voxels. This “SAC contact fraction” has a numerical value with the following interpretation. If a portion of a SAC dendrite intermingles with the arbor of a GC arbor that has SAC contact fraction *f*, on average the dendrite portion will devote a fraction *f* of its surface area to contact with the GC.

For each contacting or collocating voxel, we recorded the direction from the corresponding SAC soma centroid to the voxel in the plane perpendicular to the light axis (Fig. S6A). Based on these directions, the voxels were divided into 8 bins equally spaced on the circle. For each bin, we computed the ratio of contacting voxels to collocating voxels. This yielded SAC contact fraction versus direction, shown in the polar plots of Fig. S6 (Fig. S6B, D). The overall preferred direction of SAC contact for a GC is computed by taking the vector mean of the polar plot for that GC.

The SAC contact fraction for a GC is a normalized rather than absolute measure of SAC contact. The normalization is intended to make the SAC contact fraction robust to incomplete sampling of SAC dendrites. Reducing the number of SACs in the analysis tends to reduce both contacting and collocating voxels by the same factor, leaving the ratio unchanged. Our sample of SAC dendrites is biased, because it contains no SAC dendrites that come from a SAC soma outside the patch. The bias is least for GCs near the center of the retinal patch, and greatest for GCs near the borders. Our normalization procedure does a good job of correcting for the biased sampling, as shown by the reproducibility of the polar plots within each type in Fig. S6.

### Directional preference of SAC contact

SACs are known to obey the “mosaic principle,” meaning that their somas are placed as if they repel each other [Whitney et al., 2008]. However, SACs violate the stronger tiling principle. Their arbors are highly overlapping, with a coverage factor greater than 30 [Keeley et al., 2007]. This means that any retinal location is near SAC dendrites pointing in all directions. We analyzed whether GC arbors “prefer” to contact SAC dendrites of certain directions, even though SAC dendrites of all directions are available. Such contact analysis turned out to be useful for separating the On-Off and On DS cells into types, as will be described below.

For each GC, we computed the number of SAC dendrite surface voxels in contact with the GC, and divided by the amount of SAC dendrite surface voxels that *could have* come in contact with the GC (judged by proximity). The ratio was computed for each of eight bins uniformly spaced on the circle, after assigning SAC dendrite surface voxels to bins based on the direction vector from the relevant SAC soma to the surface voxel (Fig. S6A). The ratio is the “SAC contact fraction” for the GC, and quantifies directional preference of SAC dendrites for the GC.

When applied to On-Off DS cells, our analysis separates them into four types (37d, 37v, 37r, 37c) with four distinct PDs (dorsal, ventral, rostral, caudal) of SAC contact (Fig. S6B). This conflicts with reports of more than four On-Off types [Rivlin-Etzion et al., 2011, Dhande et al., 2013], but it would be difficult to further divide any of our four types without violating the density conservation principle. The average stratification profile of one type (37d) differs slightly from the other three in the relative heights of the On and Off peaks (Fig. S6C). However, due to within-type variability, the On-Off ratio cannot be used to accurately distinguish 37d from other types on a cell-by-cell basis (data not shown). This is consistent with previous failures to distinguish between On-Off DS types based on properties of their dendritic arbors [Oyster et al., 1993].

Our analysis also separates On DS cells into four types with four distinct PDs of SAC contact (Fig. S6D). One type has a unique average stratification profile: 7o has more mass in the central sublaminae than the three 7i types (Fig. S6E). This property can be used to reliably distinguish 7o cells (data not shown), but we have found no other way to distinguish between the three 7i types besides the SAC contact analysis. Our 7o corresponds with the nonclassical transient On DS type described by Baden et al., [2016] and Sabbah et al. [2017].

For the On-Off DS types, it is known that the PD of SAC contact is anti-parallel to the PD for visual motion on the retina [Briggman et al., 2011]. Equivalently, the PD of SAC contact is parallel to the PD for visual motion in the world, given the inversion of the retinal image by the optics of the eye. Therefore, our anatomical types 37d, 37v, 37r, and 37c are expected to correspond to physiological types that prefer motion in the superior, inferior, anterior, and posterior directions in the world, respectively.

Our analysis reproduces the main finding in the original publication on the e2198 dataset [Briggman et al., 2011], with two added novelties. First, we determine the PD of SAC contact by an analysis that includes all contact. Briggman et al. [2011] included only varicose contacts, because these were found to identify synaptic contacts with high accuracy. Our directional bias is weaker, because our analysis includes incidental as well as synaptic contacts. Second, we have performed the SAC contact analysis for all anatomically defined On-Off DS cells in e2198, including some cells without observable calcium signals. This enabled us to reconstruct complete tilings for all four types, and verify that every cardinal direction corresponds to a single On-Off type.

### Arbor density, complexity, and asymmetry

We projected the voxels of each cell’s volumetric reconstruction onto the 2D plane orthogonal to the light axis, and found the convex hull of this projection (Fig. S5I). The arbor density was then computed as the length of arbors divided by the area of the convex hull. The length of arbors is calculated from the number of skeleton voxels scaled by the correction factor, which is the average ratio of arbor length and the number of skeleton nodes of 381 cells. For each skeleton, a branch point was defined as a voxel with degree greater than or equal to 3. Path length was defined as the sum over all edge lengths in the skeleton, where edge length is the Euclidean distance (corrected for voxel anisotropy) between the pair of nodes joined by an edge. Arbor complexity was defined as the number of branch points in a skeleton divided by its path length.

The arbor vector was defined as pointing from the soma centroid to the skeleton centroid, after projection onto the 2D plane orthogonal to the light axis and scaling the y and z coordinates to match in physical length (Item S4J). For each type, we computed the mean of arbor vectors over the cells in the type. To quantify asymmetry for an individual cell, we constructed a plane through the soma centroid that was perpendicular to the mean arbor vector for its type. An index of asymmetry was defined as

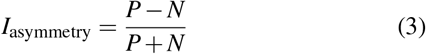

where *P* is the amount of skeleton on the same side of the plane as the mean arbor vector, and *N* is the amount of skeleton on the other side. Statistical significance of arbor asymmetry for a type was assessed by applying the Rayleigh z-test [Brazier, 1994] to the arbor vectors. The resulting *p* values were evaluated under a Bonferroni correction (Item S4N). Note that arbor asymmetry was only computed after the clustering was complete; it was not used to create the clusters.

In addition to stratification profile, soma size, and SAC contact, our hierarchical clustering also made use of two morphological properties extracted from the skeleton representations. We defined “arbor density” as the total path length of the skeleton divided by the area of the convex hull of the 2D projection of the cell onto the plane orthogonal to the light axis (Fig. S5I). We defined “arbor complexity” as the number of branch points in the skeleton divided by the total path length (Fig. S5I). Our clustering procedure used arbor density to distinguish 1ws from 1no and 1ni (Item S1, split e-5), and 2i from 2aw (Item S1, split e-8). Arbor complexity was used to separate 51 from 5ti (Item S1, split c-3), 25 and 2an from 2aw, 2i, 3o, 3i (Item S1, split e-4), and 2o from 1wt (Item S1, split e-11).

Arbor density and complexity can also be used to validate clusters that were defined otherwise, increasing confidence in the clusters. For example, it is easy to distinguish 1ws from 1ni (Fig. S5J) or 5ti from 5to (Fig. S5K) using either arbor density or complexity (Item S4H, I). These are alternatives to our clustering procedure, which used small but reproducible differences in stratification profile to make the same distinctions.

Of all types, 1ws has the lowest median arbor density (Item S4H). This extreme sparseness is consistent with our identification of 1ws as the M1 melanopsin cell [Berson et al., 2010]. 1ws had a weak and noisy light-evoked response (Fig. 4A, Fig. S7A), consistent with reports that M1 cells, though intrinsically photosensitive, have sluggish responses to ambient light [Berson et al., 2002]. Likely correspondences with other melanopsin types are 9w to M2 and 8w to M4 [Estevez et al., 2012].

Of all types, 5ti has the highest median arbor complexity (Item S4I). Helmstaedter et al. [2013] previously identified 5ti and 51 as two candidates for the W3 cell, which was originally defined and studied in a transgenic mouse line [Zhang et al., 2012]. Both types have their primary arbor at the On-Off boundary of the IPL and also have some dendrites extending as far as the INL. Our most likely W3 candidate is 51, because its secondary arbor is near the INL, and it responds omnidirectionally to both On and Off edges of the moving bar [Zhang et al., 2012, Sümbül et al., 2014]. We regard 5ti as a less likely W3 candidate because its secondary arbor is peaked near the Off SACs rather than the INL, and it does not consistently display omnidirectional responses to the moving bar. On the other hand, Zhang et al., [2012] claimed that W3 is the most numerous type in the mouse retina, while 5ti is most numerous in our sample (Fig. 4C).

A further morphological property that we studied was asymmetry of the dendritic arbor, motivated by the fact that five asymmetric types have been identified by genetic means [Kim et al., 2008, Rousso et al., 2016]. For most cells of the J type, defined by expression of JAM-B, the dendritic arbor is asymmetrically placed on the ventral side of the soma [Kim et al., 2008]. The F family, defined by expression of Foxp2, contains four asymmetric types [Rousso et al., 2016]. F-mini^Off^ and F-mini^On^ are smaller and more numerous than F-midi^Off^ and F-midi^On^. All but one of the types have ventrally directed arbors. The exception is F-mini^On^, which is dorsally directed in the ventral retina, and ventrally directed in the dorsal retina. J and F-mini cells exhibit weak DS with a ventral PD on the retina, while F-midi cells are not DS.

We did not use arbor asymmetry during our clustering procedure. This would likely have been unreliable for separating cells into types; asymmetry is a noisy property because many arbors are cut off by the boundaries of the imaged volume. We found it useful to evaluate arbor asymmetry after clusters had already been defined. For each cell, we computed an “arbor vector” with tail at the centroid of the soma and head at the centroid of the skeletonized arbor (Item S4J). We characterized the asymmetry of each type by examining its arbor vectors for a bias towards one direction (Item S4M).

2an (Item S4K), 6t, and 63 are types that correspond with F-mini^Off^, F-midi^On^, and F-mini^On^, respectively. This correspondence is supported by comparison with data of Rousso et al. [2016] on stratification profile, areal density of cell bodies, coverage factor, and arbor asymmetry. The 63 arbor typically points in the dorsal direction (Eyewire Museum), consistent with the location of the e2198 retinal patch being in the ventral retina [Rousso et al., 2016]. 63 also corresponds to gc31-56 previously reported by Helmstaedter et al. [2013]

2aw (Item S4L) has a stratification profile consistent with both F-midi^Off^ and J. 2aw is likely a mixture of both types, because it contains three pairs of colliding or nearly colliding cell bodies (Eyewire Museum), and such collisions are forbidden in a pure type by the mosaic principle. The coverage factor is large (Fig. 4D), consistent with the reported coverage factors of roughly 6 and 1 for J and F-midi^Off^ [Rousso et al., 2016]. It is surprising that we cannot distinguish between J and F-midi^Off^, because J cells are typically much more asymmetric [Rousso et al., 2016]. However, no cell in our entire population exhibits the strong asymmetry that is typical of the J cell. One possible explanation is that a minority of J cells has symmetric arbors, and this J variety is found at the dorsal and ventral margins of the retina [Kim et al., 2008]. Perhaps the e2198 retinal patch comes from a transitional zone between asymmetric and symmetric varieties of J cell.

### Coverage factor

Each cell’s dendritic field was approximated as the 2D convex hull of its volumetric reconstruction (Fig. S5I). The coverage factor for a cluster was computed as the sum of cells’ hull areas divided by the area of the union of hulls. Border effects will lead to underestimation of the coverage factor, similar to the above-mentioned underestimation of the arbor density. To minimize border effects, we considered only the intersection of each convex hull with the crop region.

### “Confirmed” types

Our 5ti, 5so, and 5si correspond to the HD family of Jacoby and Schwartz [2017]. They co-stratify with 51 (W3b), but can be distinguished using small differences in stratification depth. 5ti (UHD) is the most numerous cluster and is less than 8% of our sample.

4i and 4on co-stratify with 4ow (transient Off alpha), but 4ow is obviously different because of its large soma size (Item S1, split b-2). The 4i stratification profile spans a larger range of IPL depths than 4on (Item S1, split b-3). The stratification profiles of 4on and 4ow are almost identical. The physiological responses of 4i and 4on are similar to those of 4ow in our dataset. One or both may correspond with a “mini alpha” type identified by Baden et al., [2016].

6sn co-stratifies with 6sw (transient On alpha), but has a smaller soma. 6sn may correspond to a “mini alpha” type identified by Baden et al., [2016].

Our 73 corresponds to the On delayed cell studied by Mani and Schwartz [2017], based on its delayed response and many recursive dendrites spanning the central sublamina.

73 is smaller than 72, and has more stuff in central sublamina. Mani and Schwartz said On delayed has more recursive dendrites amd relatively small dendritic fields.

Our 82wi corresponds to the vOS cell studied by Nath and Schwartz [2016], which is activated by a vertically oriented stimulus. The stratification profile matches, and the 82wi cells in our sample with calcium imaging data also exhibit orientation selectivity (Eyewire Museum).

### Helmstaedter et al. (2013)

Five of 12 GC clusters in Helmstaedter et al. [2013] can be placed in one-to-one correspondence with ours (Table S1). Their gc10-40 corresponds to our 25, a bistratified cell with a small secondary arbor right at the On-Off boundary. Their gc37-46 corresponds to 4i. Their gc31-56 corresponds to 63, the only central type that stratifies all the way from the Off SAC to the On SAC. Their gc36-51 and gc44-52 correspond to our 5ti and 51, respectively. 51 has a small secondary arbor close to the INL, while the 5ti secondary arbor is “taller.”

The other 7 GC clusters of Helmstaedter et al. [2013] are ambiguous and appear to contain more than one of our clusters. gc14-30 could contain 2aw, 2i, and 2o. Their gc15-42 could correspond to 2an or 1wt. gc21-69 could contain 72 and 73. gc30-63 contains 37d, 37v, 37r, and 37c. Their gc35-41 could correspond to 4on or 4ow. gc47-57 could contain 5so and 5si. gc76-86 could contain any of our clusters starting with “8” and “9.”

Only 12 GC types were identified mainly because the sample is so small (45 GCs) that it contains only zero or one example of less common types. For comparison, note that the present volume (e2198) is about (0.3 mm)^2^, and contains almost 400 GCs.

### Prevalence

Our estimates of prevalence are typically lower than those reported in the literature. For example, the W3 cell has been estimated at 13% [Zhang et al., 2012] and 8% [Sanes and Masland, 2015] of the total GC population. Our corresponding 51 type is less than 5% of our GC sample.

Sanes and Masland [2015] defined a list of “securely known” types that is a subset of ours. (Our list of securely known types also includes all F types in Rousso et al., 2016.) They estimated that their list encompasses 60% of all ganglion cells, and concluded that the task of ganglion cell classification is over half done. It turns out that their list amounts to only 32% of our sample (Fig. 4C). Sanes and Masland [2015] further guessed “that there are around 30 types that compose at least 1% of the population each (there could be any number of extremely rare types) and that together they account for ≥95% of all RGCs.” In fact, our 35 validated types together account for only 84% of our sample.

### Density conservation test

Before quantifying density conservation, dendritic trunks were removed by eliminating skeleton nodes with IPL depth greater than the peak of the stratification profile plus 0.1.

Our sample is missing those GC dendrites inside the e2198 volume that come from cell bodies lying outside the volume. The missing dendrites lead to underestimation of aggregate arbor density. To minimize this border effect, we cropped away 1000 pixels (65 μm) on all four sides of the e2198 patch (Fig. 5D). The remaining “crop region” was divided into unit grid squares of 600 × 600 pixels, with physical dimensions of (40 μm)^2^. For each cluster, the aggregate arbor density was computed for each grid square (Fig. 5E, H). To quantify density conservation for a cluster, we computed the coefficient of variation (CV), defined as the standard deviation divided by the mean, for the densities of the grid squares. A small CV meant that the aggregate arbor density was approximately uniform.

To quantify statistical significance for each cluster, we needed to compare with randomized configurations of the arbors in the cluster. Naively we would have done this by randomly relocating every cell’s soma to a different location within the patch. However, this naive randomization would have typically decreased the amount of arbor within the patch, by causing some full branches to extend outside the patch, and some cutoff branches to terminate in the interior. Therefore, we constrained the soma relocation to approximately preserve the total amount of dendrites within the patch. For each soma, we drew a rectangular “orbit” that preserved the distance of the soma from the border of the patch (Fig. 5G). Each cell’s soma was randomly relocated to another position on its orbit. The arbor was also rotated by some multiple of 90° so that the cutoff branches remained oriented towards the border of the patch (Fig. 5G).

The true CV was compared with the CVs of 10,000 randomized configurations. A cluster was judged to have passed the significance test if its CV was less than 99% of randomized configurations.

### Analysis of calcium imaging data

Imaging was performed in a single focal plane for each tile in a 3 × 3 array of tiles. For each tile, the location of the moving bar was adjusted so that it was centered on the tile. Then the responses of all cells in the tile were recorded for this common stimulus. The bar’s speed, width, and location in the receptive field could not be tuned on a cell-by-cell basis to optimize visual responses, as is commonly done with intracellular recordings.

#### ROI extraction and detrending

Ellipsoidal ROIs were manually drawn around fluorescent cells (Fig. 1C). All pixels in an ROI were summed to yield fluorescence versus time for that cell. We converted to fractional fluorescent change (Δ*F*/*F*) relative to the mean computed over the entire recording. We then detrended the fractional fluorescence change by subtracting a Gaussian smoothed version of itself with σ = 20s, to reduce slow drifts in the baseline.

Correspondence between ROIs in calcium imaging (Fig. 1C) and cells reconstructed in EM (Fig. 1B) were established as follows: First, starting from the somata of On-Off DS cells from our classification, two experts visually examined their surroundings in the ganglion cell layer in the EM volume. Using the vasculature in the images as the guiding landmark and the shapes and arrangements of neighboring cells as the deciding features, 25 corresponding ROIs were identified in the fluorescent image. EM coordinates were recorded for roughly the center point of each of these somata, and fluorescent image coordinates were recorded for the center of the corresponding ROIs. A linear transformation between these two sets of coordinates is then computed in the least squares sense from their respective 3D or 2D homogeneous coordinates. Using this transformation, we computed the “nominal” EM coordinates for the center point of each of the ROIs. Finally, a human expert examined each of these EM locations to identify the correct cell that corresponds to the original ROI, similar as in the initial step.

#### Response quality index

To quantify reproducibility of visual responses, we used the quality index of Baden et al., 2016, defined as

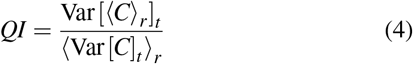

where *C* is a *T × R* “response matrix” and 〈〉*_x_* and Var []*_x_* denote the mean and variance across the indicated dimension, respectively. The number of time steps per trial is *T* and the number of trial repetitions is *R*. Cells were included in the sustainedness index calculations of Fig. 7C, E and Fig. S7B only if the quality index exceeded a threshold of 0.5.

Our recording consisted of 5 trials. Each trial contained 8 stimuli corresponding to 8 directions of motion presented in the same order (Figs. 1D, E). The recording was continuous; there was no gap between consecutive trials or stimuli. For all analyses we discard the first trial, which exhibited adaptation to the stimulus, and use only the remaining *R =* 4 trials for the response matrix *C*.

Each of the eight stimuli in a trial lasted for 31 time steps. For computing the quality index of On responses, the response matrix omitted all but the 4th to 15th time steps of each stimulus, so that each trial had *T =* 96 time steps (12 time steps by 8 directions). For computing the quality index of Off responses, the response matrix omitted all but the 15th to 31st time steps, so that each trial had *T =* 136 time steps (17 time steps by 8 directions). For computing the quality index of general responses not specific to On or Off (used by Fig. 5C, F with the same threshold of 0.5), the response matrix contained all time steps, so that each trial had *T =* 248 time steps (31 time steps by 8 directions).

#### Sustainedness index

To compute the sustainedness index of a cell in Fig. 7, we define a reference time at 0.8 sec after the nominal stimulus onset (just before stimulus offset). For the On response, we normalize the temporal response function such that the minimum is 0 and the maximum at or before the reference time is 1, and then we define the On sustainedness index as the value at the reference time. For the Off response, we normalize the temporal response function such that the minimum is 0 and the maximum after the reference time is 1. We similarly define the Off sustainedness index as the value at 0.8 sec after the nominal stimulus offset time. For inner-outer central and inner cell clusters, we computed the On sustainedness index; for outer cell clusters, we computed the Off sustainedness index (Fig. 7E).

A cell with a larger dendritic arbor might spuriously appear more sustained if that means the stimulus exits the receptive field later. To compensate for this effect, we estimate the additional time taken by the leading or trailing edge of the moving bar to exit the dendritic field. For a given low-level cluster in our hierarchy, we find the cell with the maximum diameter in the cluster, and compute the time for the leading or trailing edge of the moving bar to exit the dendritic field as half of the arbor diameter divided by the speed of the moving bar. To the “exit time” for that cluster we add 0.6 seconds, which yields a cluster-specific reference time. The cluster-specific reference times vary between 0.6 to 0.8 seconds after the nominal stimulus onset or offset time. For each cluster, we computed the sustainedness index as before, but using the cluster-specific reference time (Fig. S7B). Linear interpolation was performed for adjusted time points falling between frames.

#### Comparison of sustainedness index for high-level clusters

One-way ANOVA (Matlab `anova1`) confirmed that the high-level clusters have different mean sustainedness indices (*p* < 0.01, df_error_ = 172, df_total_ = 176) (also see Fig. 7C). Post hoc pairwise comparison between groups were performed with Tukey’s honest significant difference criterion (Matlab `multcompare`).

#### Cluster-average temporal response functions

Temporal response functions (as defined in Fig. 1E) of individual cells were first normalized in the same way as when computing their sustainedness index, and then averaged to produce cluster averages (Figs. 6C, F, 7A, D, and Fig. S7B). When standard deviation or standard error is computed and shown in a figure (Figs. 6C, F, 7A, D), the cluster average is rescaled again to have a minimum of 0 and maximum of 1 (which could have already been the case if cells in a cluster are very consitent with each other in their responses), and the standard deviation and standard error shown are scaled using the same scaling factor.

#### Dependence of visual response on direction of motion

Directional tuning curves (Item S2 and Museum) are computed as follows. Our recording consists of 5 trials with the stimulus bar moving in 8 directions in each trial in the same order (Fig. 1D, E). The recording was continuous and there was no gap between consecutive trials or between stimulus directions. We discard the first trial and average the remaining 4 trials to reduce noise.

When the moving bar enters or leaves the receptive field of a cell, the typical calcium trace usually rises quickly and then decays slowly. We model this behavior with a difference of two exponential functions.

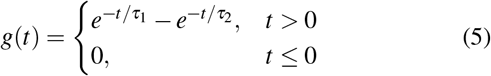

We model the entire trial-averaged response as a summation of the 8 pairs of stimulus onset and offset responses.

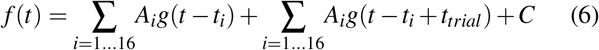

The stimulus onset and offset times *t_i_* are adjustable parameters for each cell. They are constrained to lie in narrow ranges centered on the nominal stimulus onset and offset times, defined when the leading and trailing edges of the bar cross the center of the imaged tile. The second term in the above equation is intended to model the decaying responses from the previous trial that are observed in the calcium signal at the beginning of each trial (*t*_*trial*_ is the total time of a trial with 8 directions).

For each ROI, we use the Levenberg-Marquardt nonlinear least squares method (Matlab `lsqcurvefit`) to find the set of *A_i_*, *t_i_*, *C*, *τ*_1_, *τ*_2_ parameters that best fit the recorded calcium trace. Using this fitted set of parameters, we compute the magnitude of response for stimulus event *i* as

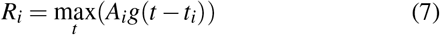

where *t* can only take values at discrete time frames corresponding to our calcium recording. For directional tuning curves, magnitudes for On and Off responses are summed together at each direction.

### Inference of the e2198 retinal patch orientation

The orientation of the compass rosette in Fig. 1A was inferred as follows. We used the average arbor vector for 2an as an estimate of the ventral direction, because 2an corresponds to a genetically defined type (F-mini^Off^) that is known to have ventrally directed dendrites [Rousso et al., 2016]. We used the preferred direction (PD) for motion of 7o to define the rostral direction on the retina (posterior direction in the world), based on the assumption that the three 7i types correspond with the classical On DS cells, which have PDs in the superior, inferior, and anterior directions in the visual world [Oyster and Barlow, 1967]. Equivalently, the PD of SAC contact of 7o defines the caudal direction on the retina, as it is anti-parallel to the PD for motion. The end result is that 2an dendrites point roughly toward 240^°^, and 7o cells have SAC contact PDs of ~170° (Fig. S6D) and physiological motion PDs of ~340°.

### Code availability

Code for the analyses of the 3D cell reconstructions and calcium imaging data will be made available on Github at https://github.com/seung-lab/e2198-gc-analysis upon publication.

### Data availability

Cell reconstructions and other data in summary forms can be viewed at http://museum.eyewire.org/. Raw calcium imaging data will be made available together with the analysing code at https://github.com/seung-lab/e2198-gc-analysis. EM volume and reconstruction data will be made available at http://seunglab.org/data/.

## Supplemental Items

### Item S1 (ItemS1.pdf)

Clustering Dendrograms and Histograms. Histograms describing each split in the dendrogram in Fig. S2A. (included in the first page; a: high-level classification, b: outer central, c: inner-outer central, d: inner central, e: outer marginal, f: inner marginal).

### Item S2 (ItemS2.pdf)

Cell Type Summary Information. Top left (a): Skeletonized dendritic tiling across the computationally flattened patch of retina. Bottom left (b): tangential view of the skeletonized dendritic tiling with the On SAC layer (lower gray dashed line) and Off SAC layer (upper gray dashed line). Note that cell bodies are artificially generated regardless of their actual sizes for both panels a and b. Top right (c): stratification profiles for each type, with the mean stratification profile shown in gray. Bottom right (d, e): directionality and mean time course of Ca response over all directions and trials.

### Item S3 (ItemS3.zip)

Physiology Summary for Individual Cells. This file contains images summarizing the physiology recordings for each cell. There are 315 files corresponding to the 315 cells with two-photon calcium imaging data, organized into 47 folders corresponding to the types in our classification. Each image shows the location of the ROI within the two-photon overview image (upper left), the calcium ΔF/F (%) during the entire stimulus delivery period of 5 trials with 8 direction each (top), averaged trial (blue) and parametric fit (red) as described in Methods (bottom), a breakdown of each trial (grey) and the averaged trial (blue) to 8 stimulus directions (middle right), and the physiology polar plot with 5x scaled version of the corresponding DS vector (lower left).

### Item S4 (ItemS4.pdf)

Morphology Summary of GC Clusters. (A) Soma size. (B) Max diameter. (C) Mean branch width. (D) Correlation of soma size and mean branch width. (E) Correlation of soma size and path length. (F) Path length. (G) Hull area. (H) Arbor density. (I) Arbor complexity. (J) Arbor vector. (K) Arbor vectors of cells in 2an. (L) Arbor vectors of cells in 2aw. (M) Arbor asymmetry index. (N) Rayleigh test p-value of arbor asymmetry.

### Item S5 (ItemS5.pdf)

Contribution of Individual Eyewirers. This file includes list of their online username within the Eyewire (http://eyewire.org) community.

### Table S1 (TableS1.pdf)

Table of Correspondence with Previous Literatures.

### Movie S1 (MovieS1.mp4)

Video Created to Celebrate the Successful Completion of Phase 3 of Countdown to Neuropia. The neurons are all those with cell bodies in a 0.2 × 0.2 mm2 patch of the GCL covered by Countdown Zones 1 through 3, and are also rendered in Item S2.

